# An ancient evolutionary origin for IL-1 cytokines as mediators of immunity

**DOI:** 10.1101/2025.07.17.662885

**Authors:** Francisco Fontenla-Iglesias, Tse-Mao Lin, Amelia G. Williams, Sabyasachi Das, Chia-Yen Lee, Jonathan P. Rast, Max D. Cooper, Katherine M. Buckley, Sebastian D. Fugmann

**Author notes:** Co-corresponding authors: J.P.R., K.M.B., and S.D.F. These authors contributed equally (Francisco Fontenla-Iglesias, Tse-Mao Lin, Amelia G. Williams). Lead contact: S.D.F.

## Abstract

Inflammation is a hallmark of immune responses. Its mechanistic underpinnings in mammals are well-defined: pro-inflammatory cytokines of the interleukin 1 (IL-1) superfamily establish and support microenvironments that promote immune cell activities. Despite a growing number of reports on inflammatory processes and components of the IL-1 signaling axis in several invertebrate lineages, orthologs of these central cytokines have not been detected outside of the jawed vertebrates. Here, protein structure prediction algorithms were applied to identify genes encoding a family of IL-1 proteins with homologs throughout Eumetazoa (termed “IL-1anc” to reflect their ancestral evolutionary origin). Using *Petromyzon marinus* (sea lamprey) and *Strongylocentrotus purpuratus* (purple sea urchin) as model systems, we demonstrate that IL-1anc proteins share important features with mammalian IL-1α/IL-1β including expression patterns, protein localization, and processing. Together, our data indicate that the IL-1 superfamily and associated circuitry represent a foundational module of animal immunity that far predates the jawed vertebrates.

## Introduction

Multicellular organisms rely on cell-to-cell communication to efficiently orchestrate responses to microorganisms. Upon detecting pathogen-associated molecular patterns (PAMPs), pattern recognition receptors (PRRs) initiate intracellular signaling cascades that culminate in the production of cytokines that amplify, shape and sustain the nascent innate immune response. This state is commonly referred to as inflammation which, in vertebrates, is regulated by the production of interleukin-1 (IL-1) family cytokines (Dinarello, 2018).

The mammalian IL-1 superfamily consists of 11 members that are broadly divided into three subfamilies (named for their founding members: IL-1, IL-18 and IL-36)(Rivers-Auty et al., 2018). The defining characteristic of IL-1 superfamily proteins is a conserved C-terminal β-trefoil fold (Krumm et al., 2014). Phylogenetic analysis of the vertebrate IL-1 family indicates that IL-1α, IL-1β, IL-36, IL-37, and IL-38 arose from a common ancestor (Rivers-Auty et al., 2018; Wang et al., 2025). Surprisingly, however, both IL-18 and IL-33 appear to be only distantly related to the rest of the IL-1 superfamily, raising the possibility that the last common ancestor predates the emergence of extant jawed vertebrates. Notably, the β-trefoil fold structure is not unique to IL-1 proteins but is also present in other protein families, including fibroblast growth factors (FGFs) (Faham et al., 1998). The IL-1 receptor system is similarly conserved: two or three extracellular immunoglobulin (Ig) domains bind IL-1 ligands and a single, intracellular Toll/Interleukin-1 Receptor (TIR) domain activates signaling (Boraschi et al., 2018). Whether the conserved structural similarity among the IL-1 ligand and receptor proteins represents convergent evolution or a deep ancestral connection remains an open question.

The best-characterized members of the IL-1 superfamily are the closely-related IL-1α and IL-1β (Dinarello, 2018). Phylogenetic analysis suggests that the genes encoding these cytokines are the result of a tandem gene duplication that occurred at the origin of mammals(Rivers-Auty et al., 2018). Subsequently, however, these paralogs have each evolved a unique character. IL-1α is constitutively expressed by a broad range of non-hematopoietic cell types including epithelial cells and keratinocytes where it localizes to the nucleus (Di Paolo and Shayakhmetov, 2016). In contrast, IL-1β is transcriptionally upregulated in myeloid cells in response to specific PAMPs and localizes to the cytoplasm as an inactive pro-form (Dinarello, 2018). The IL-1β protein is activated when innate immune signals trigger the assembly of inflammasomes (complexes of Nod-like receptors (NLRs), pyrin and caspase-1) resulting in the activation of caspase-1, a protease that cleaves two substrates: pro-IL-1β and gasdermin D (GSDMD)(Schroder and Tschopp, 2010). The N-terminal GSDMD domains multimerize to form pores in the plasma membrane through which mature IL-1β and other factors are released. This breakdown of membrane integrity ultimately results in pyroptosis, a proinflammatory form of programmed cell death (Brodsky and Monack, 2009; Schroder and Tschopp, 2010).

Both IL-1α and IL-1β lack signal peptides and are secreted via non-classical pathways. The cleavage of IL-1β is essential for its secretion and biological activity. The mechanism by which IL-1α is secreted remains unclear, although the IL-1α pro-peptide can be processed into a mature form by several proteases including calpain (Dinarello, 2018). IL-1α can also be retained on the outside of the cell via interactions with lectins (Kurt-Jones et al., 1985; Kaplanski et al., 1994), whereas IL-1β exists only in a free, non-membrane-bound state in the extracellular space. Despite these mechanistic differences, both IL-1α and IL-1β signal through the same IL-1 receptor (IL-1R) complexes (Boraschi et al., 2018). Ligand binding to IL-1R triggers autocrine and paracrine signals that act in a feed-forward loop to induce the production of additional pro-inflammatory molecules (Boraschi et al., 2018).

Many paradigms established in mammalian innate immune systems have been broadly applied across other classes of jawed vertebrates, as well as the jawless vertebrates and invertebrates that lack V(D)J-based adaptive immune systems. Pro-inflammatory cytokines remain as a singular exception to this rule. Homologs of only one such family of mammalian cytokines (IL-17) have been identified and implicated in invertebrate immune responses (Roberts et al., 2008; Wu et al., 2013; Buckley et al., 2017). While attempts at identifying IL-1 homologs based on sequence similarity have not yielded any candidates, potential invertebrate IL-1Rs have been readily identified based on conserved domain structure (Hibino et al., 2006), although these domain organizations may be convergent (Zhang et al., 2010). Furthermore, components of the inflammasome pathways regulating IL-1β activation and secretion are present throughout invertebrates, including NLRs (Yuen et al., 2014; Buckley and Rast, 2015; Degnan, 2015), caspases (Robertson et al., 2006; Xu et al., 2011), and gasdermins (Jiang et al., 2020; Qin et al., 2024). Finally, the recent discovery of pyroptosis-related programmed cell death pathway in corals, hydra and amphioxus that rely on caspase-mediate gasdermin cleavage provides the first experimental evidence that this aspect of inflammation is phylogenetically widespread (Jiang et al., 2020; Chen et al., 2023; Wang et al., 2023). This raises the possibility that the IL-1 superfamily may also be functionally conserved within Metazoa.

The above findings collectively suggest that orthologs of the pro-inflammatory IL-1 cytokines may exist in invertebrates. One possibility why they have not been discovered is related to the fact that co-evolution with pathogens drives rapid diversification in metazoan genes involved in the immune response (Vinkler et al., 2023). Moreover, cytokines are known to be particularly poorly conserved in terms of their primary amino acid sequences even among jawed vertebrates (Li et al., 2007; Dijkstra, 2014), and the relatively small size and complex exon/intron structure of IL-1 superfamily members (*i.e.,* IL-1 domains are not typically encoded in single exons) complicate the identification of distant orthologs of this gene family. Although sequence identity among IL-1 orthologs may be too low to detect across broad phylogenetic clades, the presence of putative IL-1Rs with broadly conserved ligand-binding and signaling domains suggests that the three-dimensional structures of their ligands may also be conserved. The recent development of high-quality protein structure prediction (Jumper et al., 2021) and related search algorithms (Van Kempen et al., 2023) has allowed us to revisit the quest for orthologs of IL-1 cytokines in invertebrates.

Here, we present evidence to support an ancient origin of the IL-1 cytokine superfamily within basal eumetazoans. By integrating predicted structural similarity and conserved micro-and macrosyntenic chromosomal regions with key molecular features and important functional similarities in the context of immune responses, we have discovered a previously uncharacterized, divergent family of IL-1 orthologs in jawed and jawless vertebrates as well as a wide range of invertebrate phyla.

## Results

### IL-1 orthologs are present throughout cartilaginous and jawless fish

To identify members of the IL-1 superfamily in jawless vertebrates, a PSI-BLAST search was performed using the amino acid sequences of IL-1β from *Homo sapiens* (human), *Oncorhynchus mykiss* (rainbow trout) and *Callorhinchus milii* (elephant shark) as queries against the reference protein database generated from sea lamprey (*Petromyzon marinus*; Suppl. Table 1). Three candidate sequences were identified as potential IL-1 homologs based on sequence similarity, termed here PmarIL-1.1, PmarIL-1.2, and PmarIL-1.3. Although the sequence similarity to the vertebrate cytokines was low (Suppl. Fig. S1), structural predictions using both Alphafold2 (Jumper et al., 2021) and ESMfold (Lin et al., 2023) followed by FoldSeek (Van Kempen et al., 2023) as well as comparisons using MatchMaker in Chimera (Meng et al., 2006) and the Pairwise Structure Alignment tool at the RCSB PDB (Bittrich et al., 2024) revealed that the proteins encoded by the three candidate *P. marinus* IL-1 genes each form the characteristic β-trefoil fold structure that typifies members of the IL-1 superfamily (Fig. 1A-D)(Krumm et al., 2014). Importantly, the sequential order of 12 β-strands along the peptide chain and the order in which they assemble into the three β-sheets that form the foils is conserved relative to mammalian IL-1 proteins (Suppl. Fig. S1). Furthermore, each of the three putative sea lamprey paralogs contains a conserved “[F/L]ES” motif in β-strand 9, which is a signature trait of many vertebrate IL-1β proteins although no function has yet been assigned to this amino acid triplet.

**Figure 1.**
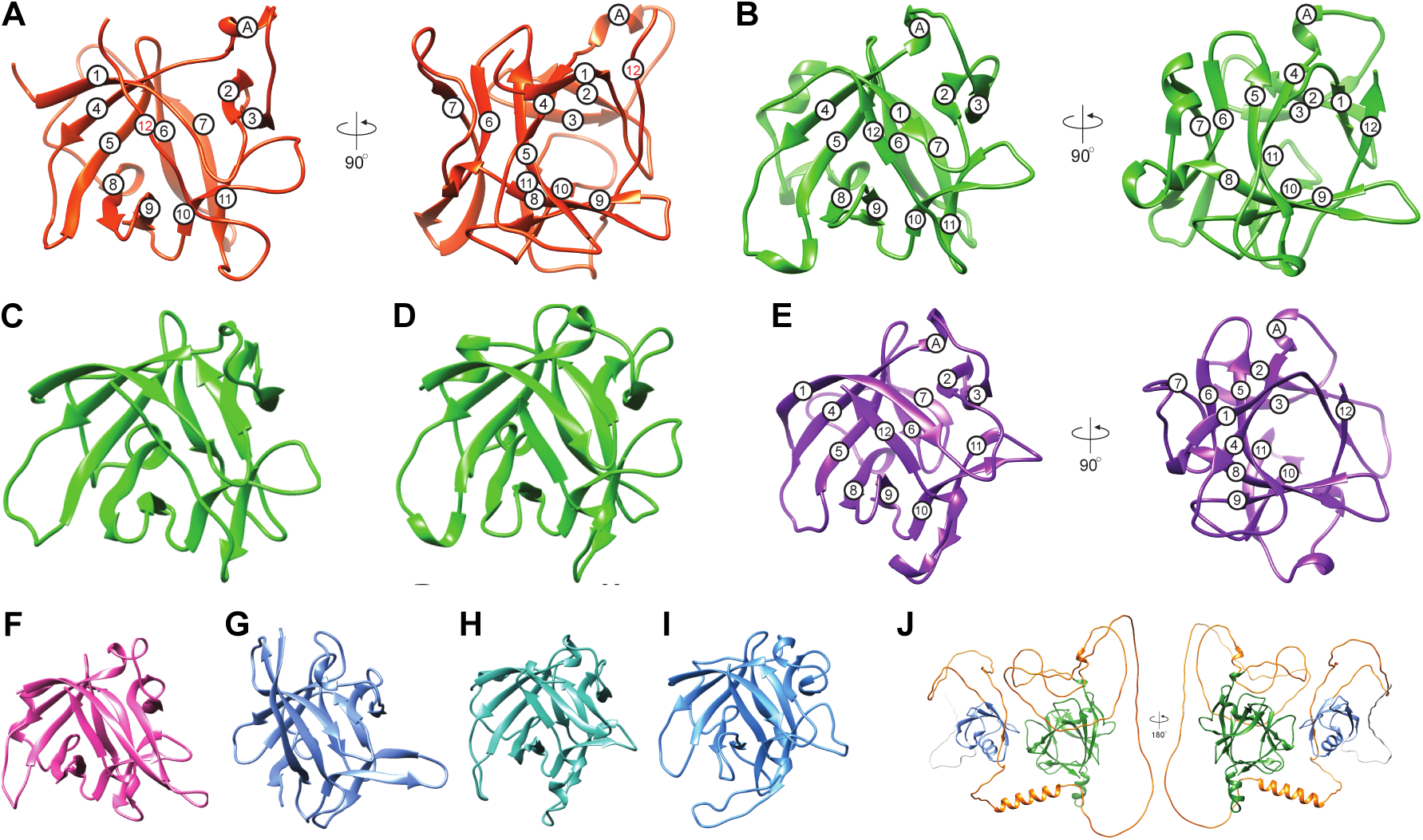
IL-1 family protein structures from divergent Metazoa. Comparison of the crystal structures of (**A**) human IL-1β (1ITB) with computationally determined structures of the IL-1 trefoil domain from (**B**) *Petromyzon marinus* (sea lamprey) PmarIL-1.3, (**C**) PmarIL-1.1, (**D**) PmarIL-1.2, (**E**) *Strongylocentrotus purpuratus* (purple sea urchin) SpurIL-1.2, (**F**) the cephalochordate *Branchiostoma floridae* BfloIL-1, (**G**) an arthropod, the crustacean *Procambarus clarkii* PclaIL-1.2, (**H**) a mollusk, the bivalve scallop *Pecten maximus* PmaxIL-1.1, and (**I**) a cnidarian, the anthozoan *Nematostella vectensis* NvecIL-1. All structures are aligned to the orientation of human IL-1β in (**A**). For (**A**, **B**, and **E**) the 12 β-strands are labeled with numbers and a small conserved α-helix region with the letter A, and they are shown from two orientations rotated 90 degrees around the vertical axis. (**J**) The predicted structure of the complete chain of *Petromyzon marinus* PmarIL-1.2 with the PDZ domain (blue) and the IL-1 trefoil domain (green) is shown from two orientations. The N-terminal region is shown in white, and the linker between the PDZ and IL-1 trefoil is shown in orange.

Analysis of the protein sequences encoded by the *PmarIL-1* candidate genes reveals that each is predicted to contain a C-terminal IL-1 trefoil domain (Pfam PF00340, and CDD cd00100 and cl00094). However, in contrast with the mammalian IL-1 superfamily proteins, PmarIL-1.2 and PmIL-1.3 are also predicted to encode an N-terminal PDZ domain (Pfam PF13180; Fig. 1J and Suppl. Fig. S2B). This is consistent with recent findings from *Cyclopterus lumpus* (lumpfish), in which a “novel” IL-1 protein (nIL-1F) also encodes PDZ and IL-1 trefoil domains (Eggestøl et al., 2020). Importantly, as with all pro-inflammatory IL-1 family members (including mammalian IL-1α and IL-1β), the PDZ domain-containing proteins from lamprey and lumpfish do not encode detectable signal peptides (Suppl. Fig. S3A) which may suggest a common, non-classical mode of secretion.

To determine if this domain architecture is widely conserved within jawed vertebrates, we searched the genome sequences of 32 non-mammalian species (11 cartilaginous, 12 teleost fish, and 7 amphibians; Suppl. Table 1) for genes with similar sequences that encode an N-terminal PDZ domain, a C-terminal IL-1 trefoil fold, and lack a signal peptide. Previously uncharacterized IL-1 superfamily members with these features (hereafter referred to as ancestral IL-1 [IL-1anc]) were identified in all chondrichthyan and osteichthyan genome sequences analyzed (Suppl. Figs. S2A and S4). Notably, analysis of the neighboring gene content indicates that, in all species, these IL-1anc genes are located in syntenic regions that are distinct from the equally-conserved region containing the classical IL-1α/β locus (Rivers-Auty et al., 2018) (Suppl. Fig. S4). The IL-1anc orthologs appear to have been lost at least twice during tetrapod evolution (at least once in the non-caecilian amphibians [Batrachia] and in a common ancestor of the crown group amniotes) and is absent from extant mammals.

The three sea lamprey IL-1 homologs are tandemly encoded within a single locus. Analysis of the neighboring genes and chromosomal gene content suggest that this locus is syntenic with the loci containing IL-1 paralogs in two hagfish species (*Eptatretus atami* and *Eptatretus burgeri*). These observations suggest that the three IL-1 orthologs arose from a series of tandem gene duplication events early within a common cyclostome ancestor (Suppl. Fig. S4). Given the independent chromosomal duplications and reorganization that shaped genomic structure in the cyclostome and jawed vertebrate lineages (Smith et al., 2018; Marlétaz et al., 2024), a lack of observed conservation of immediate neighboring genes compared to the jawed vertebrates is not surprising given the divergence time between these groups (Suppl. Fig. S4). However, the presence of numerous homologous genes within the surrounding genomic region indicate broadly conserved macrosynteny between sea lampreys and jawed vertebrates (Suppl. Fig. S5).

Together our analyses of vertebrate genome sequences reveal that: 1) IL-1 cytokines predate the origin of jawed vertebrates; 2) at least one of these ancestral IL-1 genes encoded an N-terminal PDZ domain; and 3) this ancestral IL-1 is absent in extant mammals and most amphibians coincident with the emergence of recognizable IL-1α in tetrapod genomes. A corollary of these findings is that the ancestral IL-1 genes may also predate the emergence of vertebrates. We therefore hypothesized that a similar strategy could be used to identify IL-1 superfamily orthologs within the genomes of extant invertebrates.

### Invertebrate genomes encode homologs of IL-1

Previous efforts using standard BLAST and PSI-BLAST searches and well as PFAM profile searches with HMMER failed to identify promising candidates for IL-1 homologs in a wide range of invertebrate genomic and transcriptome datasets. As an orthogonal approach, here we used the PDZ domain (PF13180) to perform HMMER searches against proteomes of representatives of the major invertebrate clades within the deuterostome superphylum: tunicates, cephalochordates, hemichordates, and echinoderms. Specifically, we focused on *Botryllus schlosseri* (star tunicate), *Branchiostoma floridae* (Florida amphioxus), *Saccoglossus kowalevskii* (acorn worm), *Asterias rubens* (common starfish) and *Strongylocentrotus purpuratus* (purple sea urchin). Protein sequences shorter than 500 amino acids were manually searched for the conserved IL-1 motifs (*e.g.,* the (F/L)ES motif) within the protein C-terminal regions to identify potential IL-1anc orthologs. Putative homologs were confirmed using structural prediction algorithms (Suppl. Fig. S6). Candidates were identified within the cephalochordate *B. floridae* (LOC118419816), the hemichordate *S. kowalevskii* (acorn worm, LOC100379003), and the echinoderms *A. rubens* (LOC117298237), and *S. purpuratus* (LOC105446168 isoforms X1 and X3, and LOC583539). Subsequent BLAST and tBLASTN searches of transcriptomes and genomes from these species and those of closely related species revealed additional IL-1 family members in these invertebrate deuterostome lineages. Potential IL-1 homologs were not identified in any of the three tunicate genomes (*Ciona intestinalis*, *Botryllus schlosseri*, and *Styela clava*) screened using this computational strategy.

A subsequent, systematic analysis of an extended range of 39 representative invertebrate genomes revealed that IL-1 superfamily proteins are encoded in many invertebrate lineages (Fig. 2, Fig. 1E-I, and Suppl. Figs. S6 and S7). It should be noted that, within each species, we numbered the predicted *IL-1* genes by the order of identification, and thus these numbers do not reflect orthology across species. There is clear evidence for lineage specific expansions in sea urchins and other echinoderms with five homologs in *S. purpuratus*. Similarly, multiple genes encoding IL-1anc proteins were identified in each species of bivalve mollusks and crustacean arthropods. Cephalochordate genomes, however, uniformly encode a single IL-1anc protein. Finally, IL-1anc orthologs could not be identified in poriferan or ctenophore genome sequences (Figs. 2 and 3). The overwhelming majority of predicted invertebrate IL-1 ortholog sequences consist of the IL-1anc domain architecture: an N-terminal PDZ domain and a C-terminal IL-1 trefoil fold, although a small subset lacks the PDZ domain (Fig. 2).

**Figure 2.**
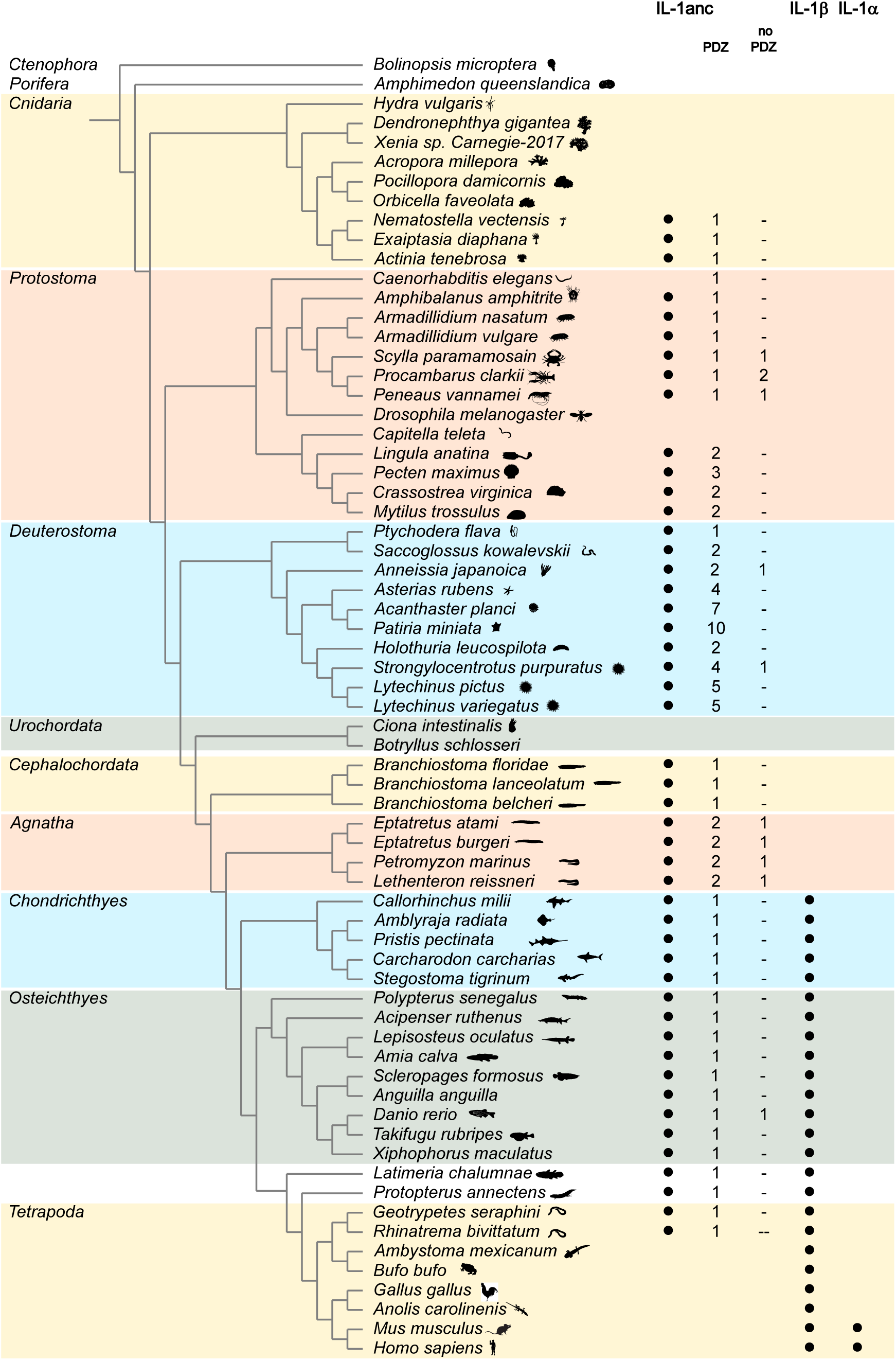
IL-1anc homologs are present throughout metazoan lineages. The distribution of IL-1 superfamily members is shown across Metazoa with representative species from major lineages. Filled circles indicate that at least one homolog is present in the respective lineage, with IL-1α and IL-1β being representative of the classical IL-1s whereas IL-1anc refers to the more widespread group described here. The numbers of predicted IL-1anc proteins with or without an N-terminal PDZ domain are listed indicating an independent loss of the PDZ domain in individual genes in various species. Note that the strategy employed in this study to identify IL-1anc proteins may have missed very divergent members of this IL-1 family in some species.

**Figure 3.**
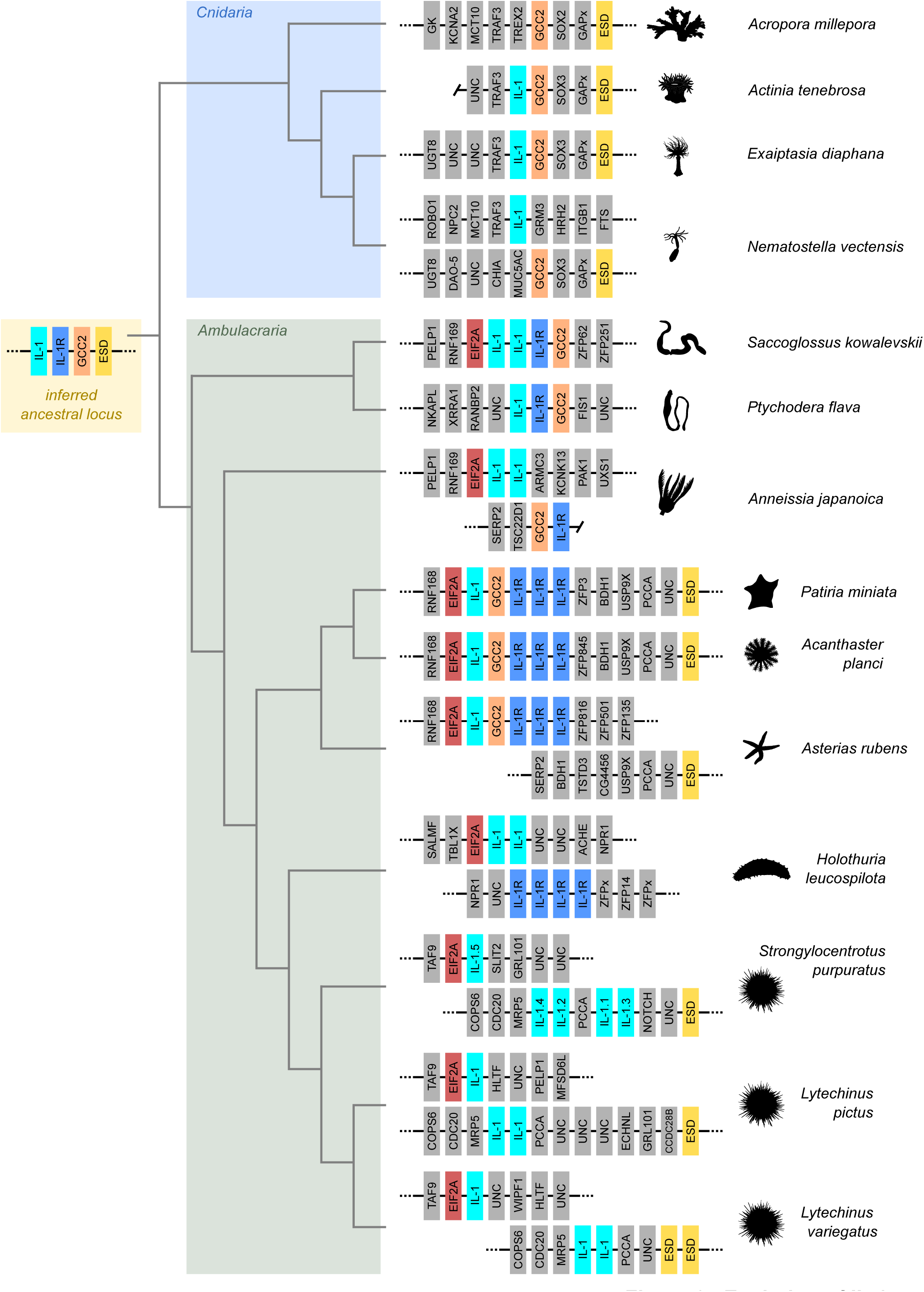
Microsynteny of *IL-1* genes in selected invertebrate phyla. A schematic representation of the chromosomal regions surrounding the most conserved *IL-1anc* family paralog (IL-1, cyan) in selected invertebrates belonging to the *Cnidaria* and *Ambulacraria* and their phylogenetic relationships. Note that additional *IL-1* genes exist outside these gene loci in several species, and that these genes were lost in corals. The IL-1s that were further characterized in subsequent experiments are numbered as they are referred to in the text instead of the generic IL-1 label. Selected genes encoding one-to-one orthologs present in multiple species are highlighted in color to emphasize the synteny between the genomic regions. An ancestral gene locus encoding both the IL-1 cytokine and its receptor (IL-1R) was inferred and is shown at the base of the phylogenetic tree.

One strong argument supporting a common ancestor of the metazoan IL-1 superfamily genes is genomic organization across divergent species. That is, conserved arrangements of neighboring genes may point to ancient homology (Zhao and Schranz, 2019). We therefore investigated the immediate chromosomal vicinity of all invertebrate *IL-1* orthologs and discovered a surprising level of conservation not only within each clade, but also among cnidarian and ambulacrarian genomes (Fig. 3). In cnidarians, *IL-1anc* genes are nearly always flanked by a homolog of *TRAF3*, and, in Hexacorallia, by homologs of genes encoding the GRIP and coiled-coil domain-containing protein 2 (*GCC2*), transcription factor Sox-3 (*SOX3*), a GTPase-activating protein (*GAPx*), and an S-formylglutathione hydrolase protein (*ESD*). Strikingly, the linkage of some *IL-1* genes to *GCC2* and *ESD* is present also in some sea stars, while hemichordates and crinoids show linkage only to *GCC2* and sea urchins only to *ESD*. All ambulacrarians share one *IL-1* linked to the eukaryotic translation initiation factor 2A gene (*EIF2A*). Surprisingly, except for sea urchins, *IL-1* genes are genomically linked to one or more gene models that encode putative IL-1R homologs (defined as transmembrane proteins with extracellular Ig folds and intracellular TIR domains). Finally, microsynteny of *IL-1* loci is conserved within cephalochordates (Suppl. Fig. S8), but does not extend beyond this clade. Taken together, an ancestral *IL-1*/*IL-1R* locus emerges from these observations in which the genes for the cytokine and its receptor are neighbored by those encoding GCC2, and S-formylglutathione hydrolase (Fig. 3).

In addition to chromosomal location, gene structures can provide insight into evolutionary conservation. We therefore assessed the exon-intron structure and phasing of representative human IL-1 family members and a set of *IL-1anc* genes from jawless vertebrates and invertebrates (Suppl. Fig. S9). In all species analyzed, the IL-1 trefoil domains are encoded by three to five exons and the phasing of two central introns in the IL1 trefoil domain is conserved and in similar positions relative to the “L/FES” motif. Additionally, when the PDZ domain is present, the two central introns in the region encoding domain are of the same phase. These observations are consistent with a shared evolutionary ancestry.

Together, these data clearly indicate that the origin of the IL-1 cytokine family precedes the emergence of vertebrates and, specifically, the vertebrate myeloid cell lineages that mediate inflammatory innate immune responses. We hypothesize that the ancestral form of IL-1 consists of a N-terminal PDZ domain linked to a C-terminal IL-1 trefoil structure and that the modern IL-1 family members in amniotes have all lost the PDZ domain.

### The ancient IL-1 proteins have immune functions

With few exceptions (Roberts et al., 2008; Buckley et al., 2017), orthologs of vertebrate interleukins have not been identified in invertebrate lineages. More broadly, invertebrate cytokines have not been well-characterized, and limited data are available on immune functions for most phyla (Watthanasurorot et al., 2011; Buckley et al., 2017; Zandawala and Gera, 2024). The discovery of IL-1 superfamily proteins throughout Metazoa thus raises the question: do ancient IL-1 proteins serve as pro-inflammatory cytokines in the context of invertebrate innate immune responses? An alternate possibility is that invertebrate IL-1 proteins act as signaling molecules in another context and were secondarily co-opted to function in the immune system coincident with the emergence of Rag1/2-mediated adaptive immunity. If the jawless vertebrate and invertebrate IL-1 proteins perform similar roles as in mammals, we anticipate that these proteins exhibit the following characteristics: 1) transcripts are primarily expressed in immunologically-relevant tissues (*e.g.*, immune cells and/or mucosal epithelia); 2) genes are transcriptionally upregulated in response to specific immune stimuli; 3) the IL-1anc proteins are cleaved and secreted in the context of a nascent immune response; and 4) interactions with an IL-1 receptor amplify innate immune responses. To investigate these properties, we conducted *in vivo* immune challenges in lamprey ammocetes and sea urchin larvae and adults, utilized publicly available transcriptome data from cnidarians, and assessed the molecular properties of jawless vertebrate and invertebrate IL-1 proteins using heterologous expression in human 293T cells.

#### Ancestral IL-1 transcripts are upregulated during immune responses

To determine if the IL-1anc transcripts are upregulated in the context of immune challenge in jawless vertebrates, lamprey ammocetes were intraperitoneally injected with either lipopolysaccharide (LPS) or flagellin. Flagellin is the primary component of bacterial flagella and a potent PAMP in many species (Zhong et al., 2017). Quantitative RT-PCR revealed that transcripts of two pro-inflammatory cytokines, *PmarTNF-α* and *PmarIL-8* (Zhu et al., 2020), were significantly upregulated 2 hours post-injection with flagellin (Fig. 4A), suggesting that the ammocetes respond to this immunological challenge. Similarly, all three lamprey *PmarIL-1 transcripts* were also upregulated in the intestine and typhlosole (a gut-associated lymphopoietic tissue) in response to flagellin but not LPS (Fig. 4A). Notably, the increase was much more dramatic for *PmarIL-1.1* and *PmarIL-1.3* (10-30 fold) than for *PmarIL-1.2*. These observations are consistent with our hypothesis that the PmarIL-1s are pro-inflammatory cytokines regulated early in infection to initiate immune responses.

**Figure 4.**
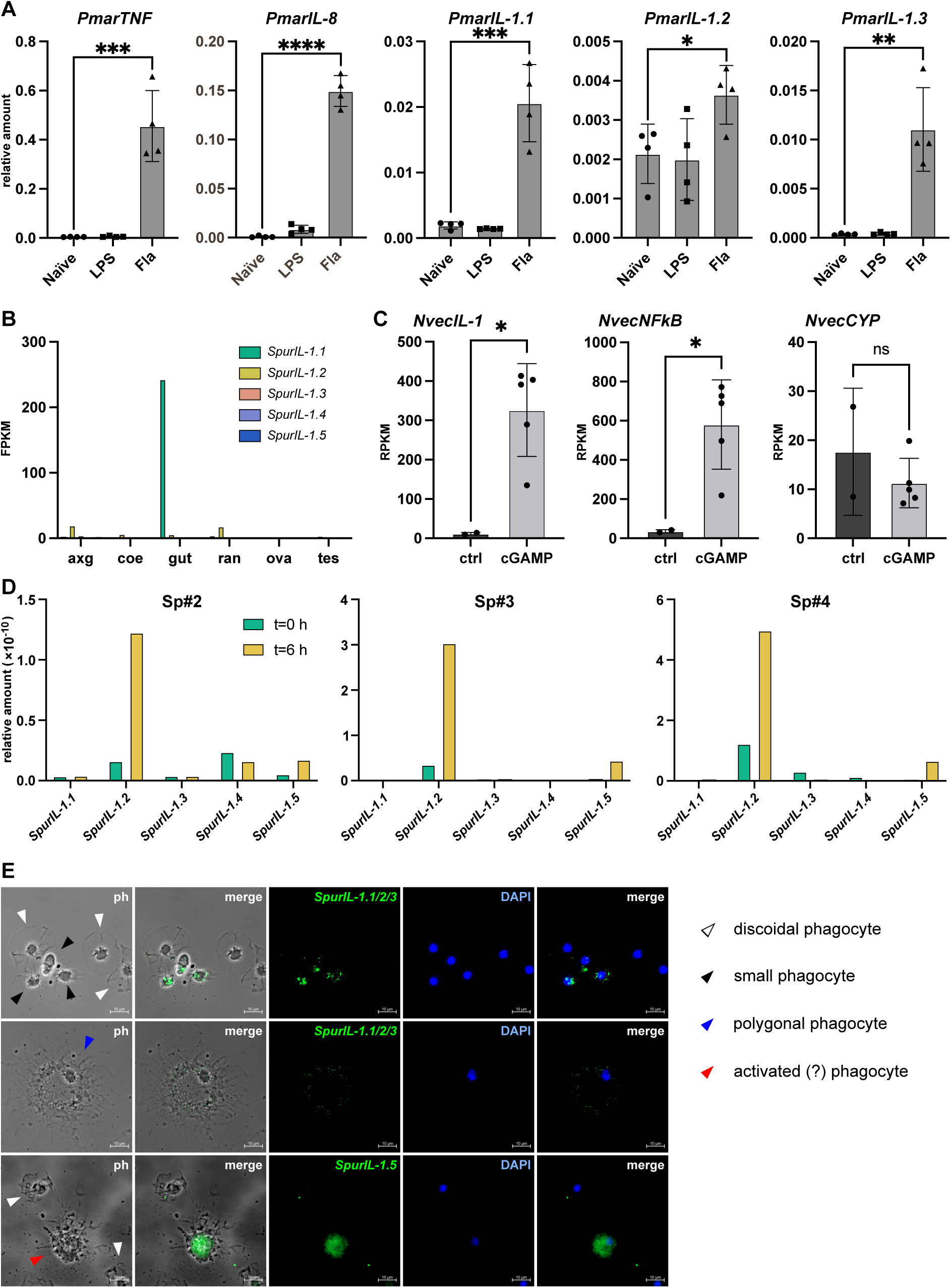
Expression of IL-1 in invertebrates and jawless vertebrates. (**A**) Three groups (N=4) of *P. marinus* ammocoetes were injected intraperitoneally with LPS or flagellin (Fla) or left untreated (naïve), and after 2 h the transcript levels of the indicated cytokines in the intestine/typhlosole were assessed by qRT-PCR. All levels were normalized to that of β-actin in the same sample and the mean ± SEM for each group is shown. A significant difference between the naïve and flagellin injected samples is indicated by asterisks (* p<0.05; ** p<0.01; *** p<0.001; **** p<0.0001, unpaired T-test). (**B**) The expression levels for each *SpurIL-1* gene in the axial gland (axg), coelomocytes (coe), gut (gut), radial nerve (ran), ovary (ova), and testis (tes) were calculated from public RNAseq datasets (see Material and Methos for details). (**C**) The expression levels of *N. vectensis IL-1anc* (LOC5506646; *NvecIL-1*), *NFκB* (LOC5504072; *NvecNFkB*), and *cytochrome p450* (LOC5501040; *NvecCYP*) in pools of untreated (ctrl, N=2) or 2’3’-cGAMP-treated (cGAMP, N=5) polyps were calculated from publicly-available RNAseq datasets (Margolis et al., 2021). A significant difference between the untreated and treated samples is indicated by asterisks (* p<0.05, unpaired T-test). (**D**) Three adult *S. purpuratus* (Sp#2, Sp#3, and Sp#4) were infected with *V. diazotrophicus*, and the expression levels of each *SpurIL-1* gene were assessed by qRT-PCR prior to infection (t=0 h) and six hours later (t=6 h). All values were normalized to the levels of *18S* transcripts in each sample, and the data for each individual is shown separately. (**E**) Fluorescence *in situ* hybridization was conducted on fixed coelomocytes (t=6 h) using the indicated HCR RNA probes with *SpurIL-1* transcripts labelled in green. The nuclei were stained with DAPI (blue), and images were collected using visual light phase contrast (ph) and fluorescence microscopy. Representative images are shown separated by channel and merged (ph+green and blue+green). The phagocytes were classified manually based on their morphology (colored arrows).

To determine whether invertebrate IL-1anc proteins also participate in invertebrate immune responses, we investigated their transcriptional regulation in larval and adult tissues of the purple sea urchin (*Strongylocentrotus purpuratus*). Five IL-1anc genes were identified using the methods described above (termed *SpurIL-1.1 – 5*). Because some the original gene models were incomplete (*SpurIL-1.3* and *SpurIL-1.5* were missing exons and *SpurIL-1.5* was spilt into separate models), we first confirmed the ORF sequence for each ortholog by cloning and sequencing expressed transcripts from adult coelomocytes. Subsequent analysis of publicly available RNAseq data from eight unchallenged adult *S. purpuratus* tissues (Tu et al., 2012) revealed that *SpurIL-1.2* is basally expressed in the axial gland, the radial nerve, and, to a lesser extent, coelomocytes (circulating immune cells), gut and testis (Fig. 4B). *SpurIL-1.1* is exclusively expressed in the gut, which is consistent with an immune function as the cells in this tissue are in continuous, intimate contact with microorganisms in the gut lumen. Expression of the *SpurIL-1.3*, *SpurIL-1.4*, and *SpurIL-1.5* genes was not detected in any adult tissues. This suggests that SpurIL-1s are regulated in a tissue-dependent manner and, at least some, may be regulated by contact with microbes.

To determine if the *SpurIL-1* transcripts are regulated in the context of an immune challenge, adult *S. purpuratus* were injected with *Vibrio diazotrophicus*. Coelomocytes were harvested prior to and 6 hours post-infection (n = 3 individuals) and transcript levels were quantified using qRT-PCR (Fig. 4D and Suppl. Fig. S10A). *SpIL-17-9* transcripts were upregulated in all three individuals, indicating an active immune response (Buckley et al., 2017). Importantly transcript levels of the *SpIL-1.2* and *SpIL-1.5* genes followed comparable patterns. In contrast *SpIL-1.1* and *SpIL-1.3* were transcripts absent or expressed at very low levels in all samples, and *SpIL-1.4* was only expressed in one individual (Sp#2), but was apparently affected by the presence of *V. diazotrophicus*. Coelomocytes are a heterogeneous population of phagocytic and secretory cell types (including vibratile cells and granular red and colorless spherule cells). HCR RNA-FISH was used to define which cell types express the *SpurIL-1* genes. Due to the sequence similarity among *SpurIL-1, −2 and −3,* transcript-specific probes could not be designed and it is therefore not possible to assign expression to a specific gene. *SpurIL-1.4*, and *SpurIL-1.5* transcripts are detectable in a few phagocytic cells at t=0 (Suppl. Fig. S11). In contrast, *SpurIL-1.1/2/3* and *SpurIL-1.5* expression was readily observed 6 hours post-infection (Fig. 4E), which is consistent with our qRT-PCR analysis (Fig. 4D). *SpurIL-1.1/2/3* expression was evident in two of the three known phagocytic cell populations (small phagocytes and large polygonal phagocytes, but not discoidal phagocytes; Fig. 4E and Suppl Fig. S11). *SpurIL-1.5* signals, however, were almost exclusively present in a distinct population of phagocytic cells with a unique morphology that has not been described previously (Fig. 4E and Suppl. Fig. S11). Together these data indicate that the *S. purpuratus* IL-1 orthologs are expressed in a tissue-specific manner, even among different immune cell populations, and that the inducible expression of *SpurIL-1.2* and *SpurIL-1.5* in phagocytic immune cells mirrors that of mammalian IL-1β in macrophages.

Exposure to *V. diazotrophicus* has also been shown to elicit a robust immune response in *S. purpuratus* larvae that includes some hallmarks of vertebrate inflammation, particularly immune cell migration (Ho et al., 2016). To determine if the *SpurIL-1* transcripts are similarly regulated in the context of larval infection, we first analyzed previously-generated RNAseq data of larvae exposed to *V. diazotrophicus* and collected at 0, 2, 4, 6, 8, 12, and 24 hours post-infection (Fig. S10B-I). Of the five *SpurIL-1* orthologs, transcripts were only detectable for *SpurIL-1.2* and *SpurIL-1.4* with *SpurIL-1.2* transcripts present in uninfected larvae and upregulated seven-fold at 6 hours post-infection (Fig. S10C) mirroring the rapid upregulation of SpIL-17-I (Fig. S10G and (Buckley et al., 2017)). In contrast, *SpurIL-1.4* was only detected at low levels at the 8 and 12 hours post-infection timepoints (Fig. S10H) which parallels the slower induction of the *185/333* immune response factors (Fig. S10 H and (Ghosh et al., 2010)). As the transcript quantification was performed on pools of whole larvae, we subsequently performed *in situ* hybridization to localize *SpurIL-1.2* expression to specific cell types and/or tissues. *SpurIL-1.2* transcripts were low or undetectable in uninfected larvae (Fig. S10J), but 6 hours post-infection, were readily observed in a ring of epithelial cells within the midgut in 23 out of 23 individual larvae (Fig. S10K). As in the sea urchin adults, this transcriptional upregulation within larvae is consistent with a role in mediating the early immune response.

Finally, to determine if the *IL-1anc* transcripts are regulated during immune responses in a basal metazoan, we re-analyzed publicly-available RNA-Seq data generated from the starlet sea anemone (*Nematostella vectensis*)(Margolis et al., 2021). Using the strategy described above, we identified a single gene encoding an IL-1anc within the *N. vectensis* genome sequence (LOC5506646; *NvecIL-1*; Fig. 3). In this species, the cyclic dinucleotide 2’3’-cGAMP (a viral mimic) acts as a PAMP that elicits broad antibacterial and antiviral immune responses. Analysis of the RNA-Seq data isolated from polyps challenged with 2’3’-cGAMP reveal that expression of *NvecIL-1* was upregulated more than 30-fold (log_2_ fold change = 5.35; p_adj_ = 4.33 × 10^−31^) compared to untreated individuals (Fig. 4C). Importantly a comparable upregulation was observed for the NFκB gene, a downstream target in innate immune signaling pathways in vertebrates (Oeckinghaus and Ghosh, 2009), but not in a unrelated housekeeping gene of the cytochrome P450 family (Fig. 4C).

Taken together, our analysis of gene expression profiles in divergent jawless vertebrates and invertebrates support the hypothesis that individual *IL-1anc* genes participate in immune responses. These transcripts are specifically expressed in immunologically-relevant cells and tissues and transcript levels are rapidly upregulated upon induction of innate immune responses. This parallels known IL-1 functions promoting inflammation in mammals and strongly suggests that the function of IL-1 superfamily members within immune systems may be ancient.

#### Subcellular location and cleavage

To characterize the sea urchin and lamprey IL-1anc proteins, complete ORFs of three *P. marinus* and five *S. purpuratus IL-1anc* transcripts were either amplified from native cDNA or synthesized for codon optimization in human cells and cloned into mammalian expression vectors harboring a C-terminal FLAG tag. To assess subcellular localization, 293T cells were transiently transfected with plasmids encoding either the full-length sea urchin or lamprey IL-1anc proteins and formaldehyde-fixed 24 h later. Expression vectors encoding human IL-1β (HsIL-1β) and untransfected cells served as positive and negative controls, respectively. Recombinant IL-1anc proteins were visualized by fluorescence microscopy using anti-V5 primary and Alexa 594-labelled secondary antibodies (Fig. 5). As expected, no signal was detected in untransfected cells (Fig. 5J); cells expressing HsIL-1β exhibited fluorescence throughout the cytoplasm (Fig. 5A), which is consistent with previous reports for mammalian IL-1β (Matsushima et al., 1986; Rubartelli et al., 1990). The sea lamprey IL-1anc proteins are generally localized throughout the cytoplasm (Fig. 5G and H). The exception is PmarIL-1.1, which is often diffusely distributed, but also appears in distinct, large clusters. This pattern was not observed for any other IL-1 in this study. An analysis of the culture supernatant of such transfectants revealed that PmarIL-1.1 is uniquely secreted (Suppl. Fig. S3B), which corresponds to the presence of a predicted signal peptide in this protein (Suppl. Fig. S3A). Therefore, one explanation for these observations is that the intracellular clusters reflect an accumulation of PmarIL-1.1 in the endoplasmic reticulum. Three *S. purpuratus* proteins (SpurIL-1.3, SpurIL-1.4, and SpurIL-1.5) exhibit cytoplasmic localization (consistent with with HsapIL-1β; Fig. 5D-F). In contrast, fluorescence corresponding to SpurIL-1.1 and SpurIL-1.2 clearly overlap with the DAPI-stained nuclei (Fig. 5B and C), which is reminiscent of the nuclear expression of mammalian IL-1α (Luheshi et al., 2009). Thus, the full-length *S. purpuratus* and *P. marinus* IL-1anc proteins (with the notable exception of PmarIL-1.1) show a localization consistent with the pro-forms of the prominent mammalian IL-1 family members.

**Figure 5.**
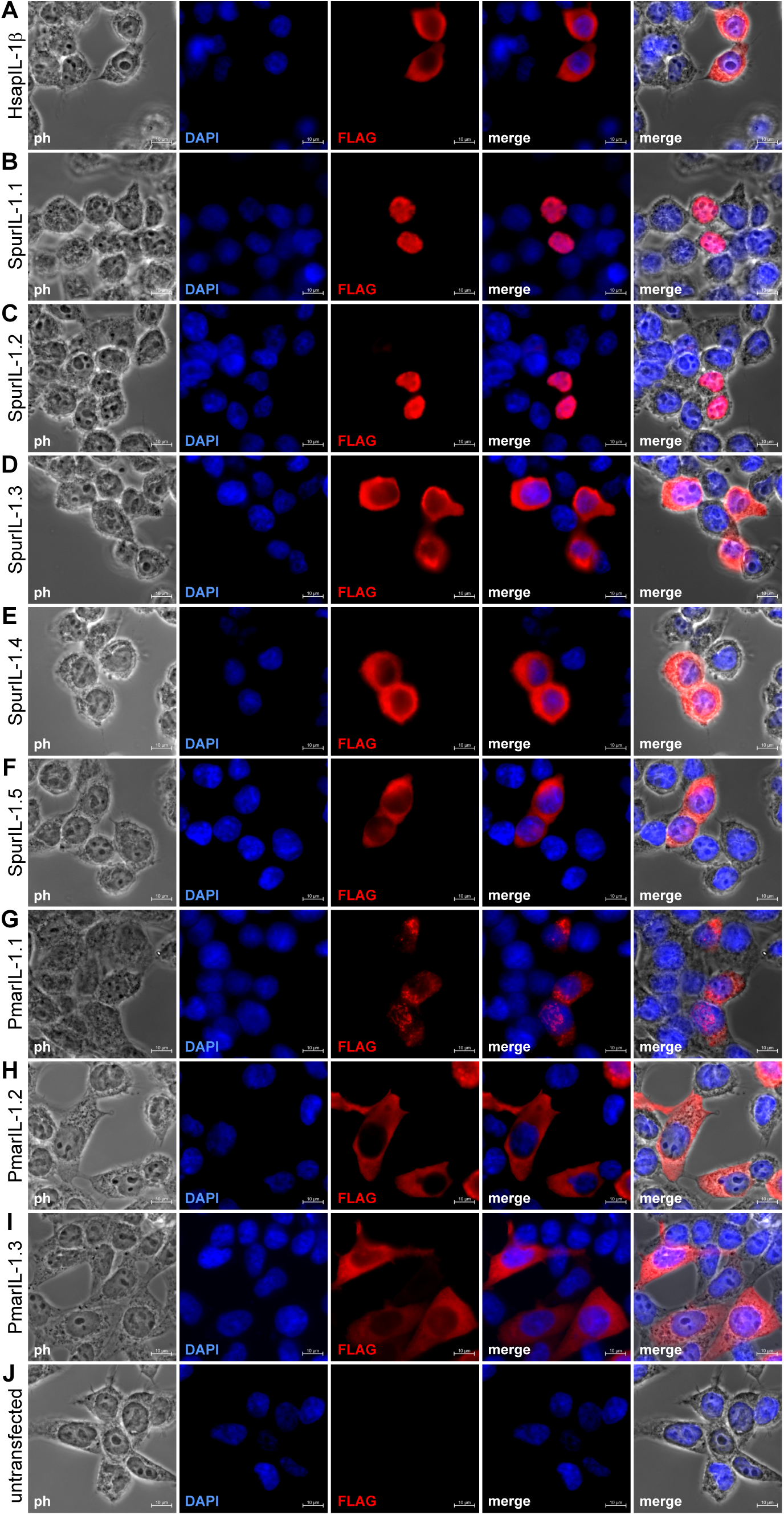
Subcellular localization of IL-1s. FLAG-tagged variants of the indicated full-length IL-1 proteins were transiently expressed in 293T cells, and their spatial distribution (shown in red) was visualized using immunofluorescence microscopy with a monoclonal anti-FLAG antibody. To classify the subcellular compartments, nuclei were stained with DAPI (blue), the cell morphology was visualized using phase contrast microscopy (ph), and images acquired in different channels were merged. Untransfected 293T cells served as negative controls.

One hallmark of mammalian IL-1β secretion is the processing of the inactive cytoplasmic form (pro-IL-1β) by caspase-1, an inflammasome-activated protease. The C-terminal fragment, IL-1β, is then released from the cell as plasma membrane integrity breaks down due to the formation of GSDMD pores (Evavold et al., 2018; Heilig et al., 2018). To test whether the *S. purpuratus* and *P. marinus* IL-1anc proteins are similarly activated by caspase activity, FLAG-tagged IL-1anc proteins were transiently expressed in 293T cells either alone or with human V5-tagged caspase-1 (HsCASP1). The sizes of the ectopically expressed proteins in the respective cell lysates were assessed by Western blot using anti-V5 and anti-FLAG antibodies. FLAG-tagged HsIL-1β served as a positive control.

Although expressed at levels lower than HsapIL-1β, in the absence of HsCASP1, all SpurIL-1 and PmarIL-1 proteins corresponded to their expected sizes (Fig. 6A and B). In contrast, in the presence of active human caspase, a 20 kD band corresponding to the mature form of HsIL-1β is clear. A similar cleaved product was observed for SpurIL-1.3 (approximately 28 kD). These C-terminal fragments were not generated when the invertebrate IL-1anc sequences were co-expressed with a catalytically inactive mutant of HsCASP1 (Δ) (Fig. 6A). We thus concluded that SpurIL-1.3 can be processed by a mammalian inflammatory caspase, albeit with far lower efficiency. Whether the other SpurIL-1 proteins are processed by other proteases (*e.g., S. purpuratus* caspases with distinct specificities) or by unknown mechanisms as in mammalian IL-1α remains to be determined. As neither PmarIL-1.1, PmarIL-1.2, nor PmarIL1.3 were cleaved by human CASP1, we also tested their processing by a putative *P. marinus* homolog of this enzyme, PmCASP1 (XP_032814408.1), but there was no evidence for a release of an N-terminal fragment corresponding to the maturation of active mammalian IL-1β (Fig. 6B).

**Figure 6.**
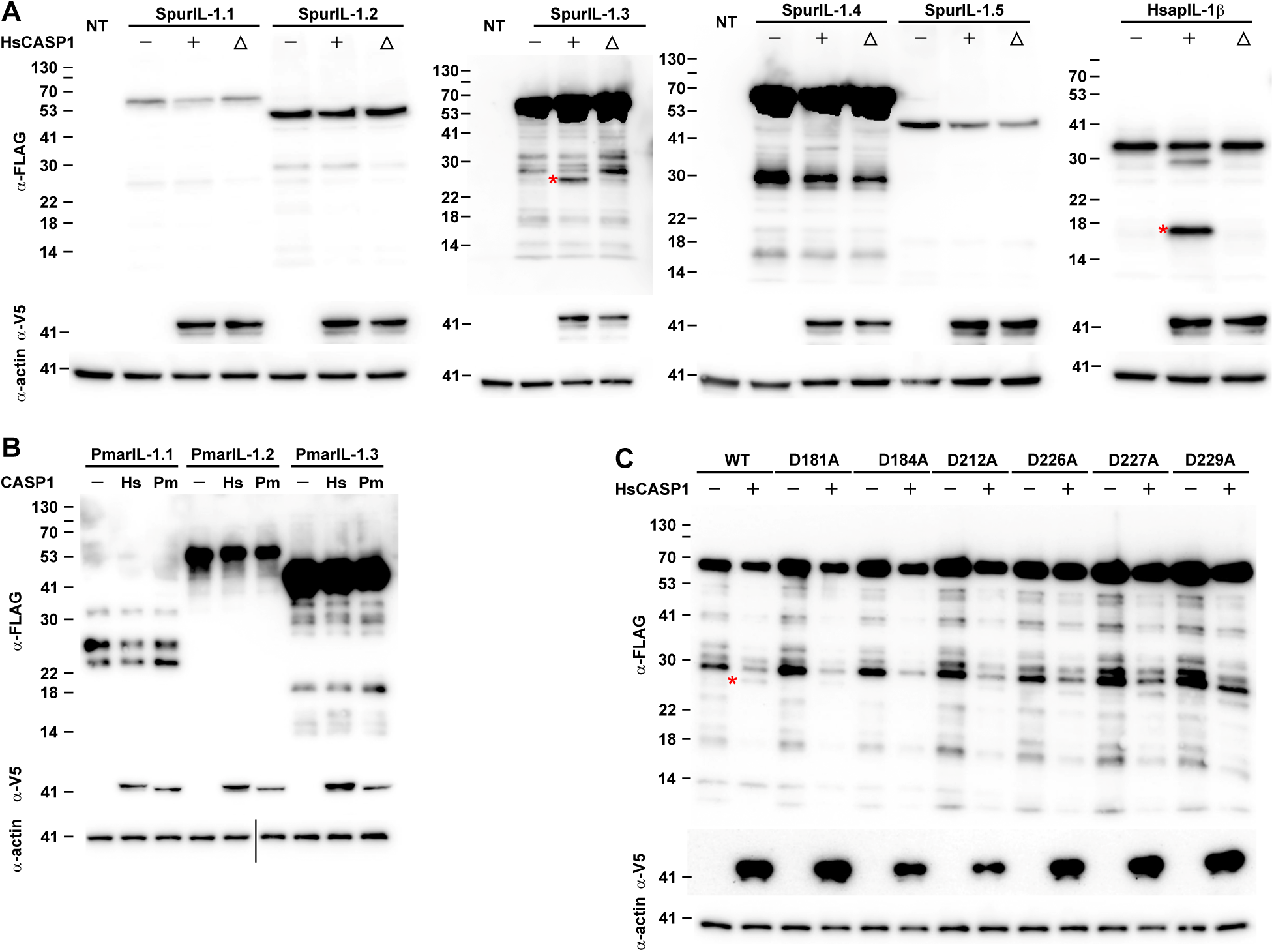
Processing of IL-1s by caspase 1. Human 293T cells were co-transfected with expression vectors for the indicated wildtype FLAG-tagged IL-1s (or mutants thereof in **C**), and V5-tagged Caspase 1 from humans (Hs, +), a catalytic mutant thereof (Δ), or *Petromyzon marinus* (Pm). Total cell lysates were resolved by SDS/PAGE and the proteins were visualized by Western blot using monoclonal anti-FLAG and anti-V5 antibodies, respectively. Non-transfected cells (NT) served as controls. The sizes of Caspase 1-dependent cleavage products are marked by a red asterisk on each blot. To illustrate that the lysates contained comparable concentrations of proteins, an anti-actin body was used. Theoretical sizes of the FLAG-tagged IL-1s are: PmarIL-1.1=33.0 kD (including signal peptide) or 26.9 kD (after cleavage of signal peptide); PmarIL-1.2=57.8 kD; PmarIL-1.3=47.9 kD; SpurIL-1.1=63.5 kD; SpurIL-1.2=55.2 kD; SpurIL-1.3=51.4 kD; mature SpurIL-1.3=30.6 kD; SpurIL-1.4=59.7 kD; SpIL-1.5=55.1 kD; HsapIL-1β=34 kD, mature HsapIL-1β-FLAG=20.6 kD.

To define the site at which SpurIL-1.3 is cleaved by HsCASP1, we individually mutated each of six putative cleavage sites to alanine. These sites were defined as aspartates upstream of the IL-1β trefoil structure (D181A, D184A, D212A, D226A, D227A, and D229A). Co-expression of these SpurIL-1.3 variants with HsCASP1 in 293T cells, revealed that only one of them was resistant to cleavage: SpurIL-1.3 D184A (Fig. 6C). This indicates that, in this system, D184 is the likely target of HsCASP1. Intriguingly the corresponding C-terminal fragment (amino acids 185-424) retains the entire IL-1 trefoil fold just as the mature HsapIL-1β resulting from cleavage of its pro-form at D116. Although the recognition motif of the cleavage site (YVHD in HsapIL-1β and DEPD in SpurIL-1.3) is not conserved, it is tempting to speculate that this cleavage of SpurIL-1.3 is an important step towards its activation.

## Discussion

Animal immune systems rely on pro-inflammatory cytokines that are rapidly activated by innate immune cells at sites of infection to initiate, sustain and amplify efficient immune responses. In mammals, the IL-1 superfamily cytokines are key players in this process. The long-standing paradigm holds that these cytokines emerged coincident with V(D)J-based adaptive immune systems at the origin of jawed vertebrates (Boraschi, 2022). The presumed absence of IL-1 orthologs in invertebrates is not due to a lack of searching, but rather, extensive searches that relied primarily on sequence similarity and predated the availability of complete genome sequences of a large number of organisms. Indeed, several studies have suggested functional equivalents based on cross-reactivity of antibodies or cytokine receptors (*e.g.,* (Beck et al., 1993; Ottaviani et al., 1993)). Here, we deployed recently developed algorithms to predict protein structures and uncover a novel addition to the IL-1 cytokine superfamily (IL-1anc) that is present not only in the majority of jawed vertebrates, but also in jawless vertebrate and invertebrate species. The *IL-1anc* genes encode similar predicted protein structures and share gene organization with the well-characterized mammalian IL-1α/IL-1β orthologs, as well as with the other members of the mammalian IL1 superfamily. Furthermore, experimental data generated during infections in three species are consistent with similar functional roles in immunity. Together, these data support a single, ancient origin of the IL-1 superfamily that includes the IL-1anc genes and dates back to a common ancestor of cnidarians and bilaterians.

### Evidence for the widespread presence of IL-1 homologs

The key to discovering the IL-1anc cytokines lies in the conserved structures of IL-1 trefoil domains, despite considerable divergence at the level of primary sequence. The structural similarities between IL-1anc proteins and IL-1 proteins from jawed vertebrates are only partially (if at all) reflected in a conservation of the primary amino acid sequence across phyla. Specifically, the short (F/L)ES motif in the ninth β strand of the β-trefoil fold is largely conserved in IL-1 proteins from cnidarians to teleosts, although its functional significance remains unknown. The distinguishing feature between “classical” IL-1 family members and the IL-1anc proteins is the presence of N-terminal PDZ domain in the latter. Importantly, an IL-1anc family member had been previously reported as a novel IL-1 in lumpfish with an unusual domain structure (Eggestøl et al., 2020). It thus unknowingly became the founding member of this widespread, evolutionarily conserved gene family.

A single origin for the genes encoding IL-1 superfamily proteins is strongly supported by genomic organization, transcriptional regulation and protein processing. Chromosomal macrosynteny is broadly conserved across phyla and microsynteny of neighboring genes is retained within groups. In some cases (*e.g.,* the cnidarian and ambulacrarian *IL-1anc* genes), microsynteny can be detected even across phyla. The intron/exon structure of most known IL-1 superfamily genes exhibit conserved intron phasing within the trefoil and PDZ domains. We here demonstrate that this conservation also extends to the jawless vertebrate and invertebrate *IL-1anc* genes. Finally, similarities within the IL-1 genes are evident within the context of an *in vivo* immune response. That is, the *IL-1anc* transcripts are significantly upregulated in phagocytic cells responding to immune stimuli, mirroring the behavior of IL-1β in mammalian myeloid cells. Specific localization of the pro-forms of IL-1anc paralogs within in cytoplasm and nucleus as well as processing by caspase-1 is also consistent with mammalian IL-1α and IL-1β. One newly discovered IL-1anc from *P. marinus* appears to be secreted via a conventional signal peptide (discussed below), which strongly suggests that the extracellular space is, in fact, the site of IL-1anc activity.

The character of mammalian IL-1α/IL-1β cytokines is complex and varies even within jawed vertebrates. While no individual invertebrate or jawless vertebrate IL-1 described here combines all the known features of IL-1α, IL-1β, or any other mammalian IL-1, it is worth noting that the diversity of this pro-inflammatory cytokine family is exemplified by the dichotomy between IL-1α and IL-1β in terms of inducibility of expression, processing, and secretion pathways. As yet there are no characters, both in terms of primary sequence and predicted structure, that clearly ally the *IL-1anc* genes with any particular member or subset of the mammalian IL-1 superfamily.

### How are ancestral IL-1 cytokines processed?

Most mammalian IL-1 proteins are cleaved by proteases; for some, this processing is essential for their secretion and the biological activity (Dinarello, 2018). Cleavage typically separates a non-conserved N-terminal sequence from the structurally conserved C-terminal IL-1 trefoil domain that serves as the ligand for IL-1R. Thus, it is conceivable that the PDZ domain of IL-1anc is removed by proteolytic cleavage to release the biologically active IL-1 domain. Obvious candidates for this protease activity include inflammatory caspases that also activated in the context of innate immune responses. Original reports of an IL-1 superfamily protein that includes an N-terminal PDZ domain did not address this issue (Eggestøl et al., 2020). In line with these expectations, our experiments that caspase 1-mediated processing cleaves the IL-1anc protein SpurIL-1.3 at a known caspase recognition site. This finding suggests two fundamental features of inflammation within animal immune systems: 1) proteolytic processing is ancient mode of regulation for pro-inflammatory cytokine activity; and 2) a two-signal requirement (the first controlling the production of a pro-form, the second the generation of the mature cytokine) is likely required to prevent inadvertent immune responses. These assumptions then raise the question of why the other IL-1anc proteins were not processed in our experimental assays? One possibility is that the cognate recognition motifs of the native inflammatory caspases differ significantly from those of human caspase 1 used here. Alternatively, as yet uncharacterized unrelated proteases may serve a similar role. Finally, it is possible that at least some these IL-1anc proteins do not require processing and are instead produced in an activated form. This scenario mirrors that of IL-1α in the mammalian immune response.

### How are the ancestral IL-1 cytokines secreted?

All previously described pro-inflammatory IL-1 cytokines lack signal peptides and are therefore secreted by unconventional (*i.e.,* ER-and Golgi-independent) secretion pathways (Dinarello, 2018). Only one of the novel IL-1anc proteins characterized here (PmarIL-1.1, which also lacks a PDZ domain) encodes a classical (albeit unusually long) signal peptide that facilitates secretion when ectopically expressed in mammalian cells (Suppl. Fig. S3A). Importantly, all hagfish genomes encode an ortholog of this protein sharing the putative long signal peptide (Suppl. Fig. S3A) indicating that this IL-1anc family member emerged in a common ancestor of all extant jawless vertebrates. Interestingly, one mammalian IL-1 family protein is also secreted in a conventional manner: the IL-1 receptor antagonist (IL-1ra)(Corradi et al., 1993). This factor binds to the extracellular domain of IL-1R1 similarly to IL-1α and IL-1β but prevents the recruitment of the IL-1R3 co-receptor, thereby blocking intracellular signaling (Schreuder et al., 1997). It is thus tempting to speculate that the IL-1.1 orthologs within jawless vertebrates serve a similar anti-inflammatory role. Although IL-1anc family members that similarly lack a PDZ domains are also present in various invertebrate lineages, these genes appear to have arisen independently (Fig. 2). Additionally, the loss of the PDZ domain in those invertebrate IL-1anc proteins is not accompanied by an acquisition of detectable signal peptides.

The pathways by which the PDZ-domain containing IL-1anc proteins might reach the extracellular space thus remains elusive. Many invertebrate genome sequences encode gasdermins and their involvement in a pyroptosis-like cell death has been shown in cnidarians and cephalochordates (Jiang et al., 2020; Wang et al., 2023) It is therefore plausible that gasdermin pores form conduits through which mature IL-1anc proteins are released similar to IL-1β secretion in mammals. Mammalian IL-1α acts as an alarmin when it is released into the extracellular space upon necrotic cell death (Kim et al., 2013). It is conceivable that at least some of the ancestral IL-1anc proteins act in a similar manner. This idea is consistent with the observation that the loss of genes encoding PDZ domain-containing IL-1s during vertebrate evolution precedes the emergence of IL-1α in mammals and raises the possibility of a handover of function between these structurally distinct IL-1 family members.

### An ancestral function for IL-1 in immune responses

Secreted mammalian IL-1 cytokines bind to transmembrane IL-1Rs expressed on the surface of a wide range of cells to activate pro-inflammatory gene expression programs. Genes encoding putative IL-1Rs are present in agnathans and have been reported in a wide range of invertebrate genomes (Hibino et al., 2006; Rast and Messier-Solek, 2008); these receptors may bind to the IL-1 ligands reported here. While the invertebrate genes differ from the jawed vertebrate IL-1R subunits that bind to IL-1 cytokines and their co-receptors in the number of encoded extracellular Ig domains (two vs. three), their intracellular TIR domains are conserved as are the downstream pathways that lead to activation of the transcription factors AP-1 and NFκB. Notably, there are also members of this receptor family with only one of two Ig domains in vertebrates, but here they serve as inactivating co-receptors rather than as the ligand-binding subunits IL-1Rs (Mariotti et al., 2023). Strikingly, in some cases, *IL-1anc* genes and putative *IL-1R* genes are genomically linked consistent with the functional interaction and coevolution of the encoded proteins. Although a role in IL-1 signaling for the invertebrate IL-1Rs still needs to be experimentally validated, it is tempting to speculate that the induction of pro-inflammatory immune genes including IL-1 and IL-1Rs themselves is an ancestral property of the IL-1 circuitry.

### What drives the rapid evolution of IL-1s?

Genes encoding cytokines, particularly the mammalian interleukins evolve rapidly (Boulay et al., 2022). This is reflected in the more direct identification of putative invertebrate IL-1Rs (Hibino et al., 2006) and our inability to reveal similarities of IL-1s across phyla using classic BLAST searches. The driving force are likely selective pressures imposed by inhibitors of IL-1 superfamily cytokines produced by pathogens with the IL-18 binding proteins encoded in the genomes all orthopox viruses being one notable example (Xiang and Moss, 1999). These inhibitors fold into a Ig domain structure that competes with the Ig folds in the extracellular domain of IL-1Rs for binding to the free cytokines (Krumm et al., 2008). As we were unable to identify IL-1 homologs in any insect species it remains possible that they diverged in terms of structure beyond recognition by our search strategies. Alternatively, a particularly efficient inhibition by some pathogens could have pushed to replace the IL-1/IL-1R axis with an alternative cytokine/cytokine receptor axis in a common insect ancestor.

In summary, our study uncovers a deeply conserved IL-1 pro-inflammatory cytokine family in jawless vertebrates and invertebrates, bridging a critical gap in our understanding of immunity outside of jawed vertebrates. Despite the challenges posed by the rapid sequence divergence of IL-1 homologs, the structural conservation, gene organization, synteny, and functional parallels between vertebrate IL-1 and its newly identified invertebrate and jawless vertebrate counterparts strongly support a shared evolutionary origin. Furthermore, these findings indicate that the IL-1/IL-1R axis has been a fundamental component of immune regulation since the emergence of Eumetazoa. Elucidating the ancestral mechanisms of IL-1 signaling not only provides new insights into the origins of vertebrate immunity but also underscores the utility of new machine learning structural prediction tools in identifying long-sought immune factors when conventional sequence-based approaches prove ineffective.

## Acknowledgements

We thank all members of the Cooper, Buckley, and Fugmann labs for helpful discussions. This work was supported by grants of the Chang Gung Memorial Hospital [CMRPD1N0312] and Chang Gung University [QZRPD176] to S.D.F., the National Science Foundation (NSF, award 2131297) to K.M.B., and the National Institutes of Health (NIH, R35GM122591 and R01AI072435) to M.D.C.

## Materials and Methods

### Bioinformatics

Initial similarity searches were performed using BLAST. PSI-BLAST was executed locally using default parameters and a variable number of iterations until convergence. Protein datasets from species of interests were retrieved from the NCBI genome repository (https://www.ncbi.nlm.nih.gov/datasets/genome/). Accession numbers for the genome sequences used in this study are shown in Suppl. Table 1. To identify protein domains of interest, HMMER v3.4^1^ was used on these datasets using the curated PDZ domain profiles (PF17817 and PF13180) obtained from InterPro (https://www.ebi.ac.uk/interpro/).

Candidate sequences were manually curated to confirm the presence of the partially conserved vertebrate F/LES motif and subsequently used as queries for additional BLAST and PSI-BLAST searches to iteratively refine the dataset. Ultimately, a combination of gene model length constraints, short regions of sequence similarity, computed structure modeling (see below), and the presence more readily identifiable PDZ domains were used to identify distant IL-1 family homologs.

The accession numbers, amino acid sequences, and common names of all IL-1anc proteins identified in this study are provided in Suppl. Table 2. When multiple *IL-1anc* genes were identified within a species, they were designated with serial decimal numbers (IL-1.1, IL-1.2, IL-1.3, etc.). These number designations are arbitrary and are not meant to imply orthology unless specifically noted.

Protein structures were predicted using AlphaFold2 (AF2) and ESMFold implemented by Google Colab with default parameters^2,3^. Structures were visualized and compared with Chimera^4^. Structures were aligned as isolated domains using Chimera MatchMaker for representation in figures and with the RCSB Pairwise Structure Alignment tool^5^ using the TM-align method^6^. Amino acid sequence alignments used to compare IL-1 domain β strands were derived from these structural comparisons. Signal peptide prediction was conducted using the PredSi online tool (http://www.predisi.de)^7^.

### Animal husbandry

#### S. purpuratus (larva)

For larval cultures, adult *S. purpuratus* were obtained from the Point Loma Marine Invertebrate Lab (Lakeside, CA) and maintained in artificial sea water (Instant Ocean; ASW) at 15 °C. Adults were fed weekly with kelp (*Macrocystis pyrifera*). Gametes were obtained through gentle shaking or intracoelomic injections of KCl (0.5 M). Embryos were grown in 0.45 μM filtered ASW in stirredvessels (30 rpm) at 15 °C, and at 4 days post-fertilization (dpf) larvae were diluted to one larva per ml ASW and fed *Rhodomonas lens* (3,000 cells/ml) every other day.

#### S. purpuratus (adult)

Adult *S. purpuratus* individuals were maintained in artificial sea water (Reef Salt, AZOO) at 14°C. All individuals used for the experiments were apparently healthy and kept in our facility for at least 1 month prior to the experiments to minimize influences from environmental changes.

#### *P. marinus* (ammocoetes)

Sea lamprey ammocoete larvae (10–15 cm in length) were purchased from local suppliers in Michigan and Maine (USA) and maintained in sand-lined aerated aquariums at 18°C. Prior to all procedures, the animals were anesthetized with buffered MS222 (Syndel) at a concentration of 1 g/L. Animals to be sacrificed were euthanized with buffered MS222 (Syndel) at a concentration of 10 g/L. All experiments were conducted in compliance with the relevant guidelines and regulations under approval from the Institutional Animal Care and Use Committee at Emory University (PROTO201700387).

### Infection models

#### S. purpuratus (larva)

Sea urchin larvae were infected with *Vibrio diazotrophicus* as described^8^. Briefly, larvae (7 dpf) were maintained at a concentration of 1 larva/ml in ASW. *V. diazotrophicus* was cultured in LB media supplemented with NaCl (600 mM final concentration; SLB) at 15°C to mid-log phase. Bacteria were collected by centrifugation (5,000 × g for 10 minutes at <u>4</u>°C) and washed three times in sterile-filtered (0.2 µM) ASW. Bacterial cell numbers were calculated based on OD_600_ absorbance values as determined by a strain-specific standard curve generated in the lab. Live bacterial counts were obtained by plating serial dilutions of the culture on SLB agar plates and counting colony numbers to calculate colony forming units (CFUs) per ml of culture. Washed bacteria were added to larval cultures at a final concentration of 10^8^ cells/ml. Larvae were collected prior to infection (t = 0h) and at specific time points after exposure (t = 3 and 6 hr) and stored in Trizol reagent (Invitrogen) at −80°C for RNA purification or fixed for HCR *in situ* hybridization.

#### S. purpuratus (adult)

Apparently healthy *S. purpuratus* individuals were infected with live *V. diazotrophicus* as previously described^9^. Briefly, 1×10^6^ bacterial cells in 150 µl sterile artificial sea water were injected into the coelomic cavity using a 1 ml syringe with a 25G×1” gauge needle. Aliquots of coelomocytes were retrieved prior to infection (t=0 h) and afterwards (t=6 h) and processed for RNA purification of HCR *in situ* hybridization (see below).

#### P. marinus ammocoetes

Four sea lamprey ammocoetes were intraperitoneally injected with either 50 ng per g of bodyweight flagellin (Sigma-Aldrich) or 5 µg per animal lipopolysaccharide (LPS of *E. coli* strain O111:B4; Sigma-Aldrich), each diluted in 20 µl of 0.66× PBS(90 mM NaCl, 18 mM KCl, 6.6 mM Na_2_HPO_4_, 12 mM KH_2_PO_4_). Uninjectedanimals servedas negative controls. After 2.5 hours, the animals were sacrificed in MS-222 according to established protocols and bled by making a cross-sectional incision anterior to the cloaca. The intestine and typhlosole— a major immune organ in lamprey larvae formed as an invagination of the intestine —were carefully dissectedand immediately snap-frozen in liquid nitrogen to preserve RNA integrity.

### RNA preparation

#### S. purpuratus (larva)

Larvae were homogenized and stored in Trizol (Invitrogen) at −80°C until analysis. RNA was isolated using the RNA Clean & Concentrator kit (Zymo Research) according to the manufacturer’s protocol and resuspended in nuclease-free water.

#### S. purpuratus (adult)

Coelomic fluid from adult *S. purpuratus* was collected through the peristomial membrane using a 1 ml syringe with a 25G×1” needle preloaded with an equal volume of calcium-and magnesium-free sea water containing EDTA and HEPES (CMFSW-EI; 449 mM NaCl, 9 mM KCl, 2.5 mM NaHCO_3_, 33 mM Na_2_SO_4_, 70 mM EDTA, 20 mM HEPES pH=7.0). The coelomocytes were pelleted by spinning at 450×g for 5 min at 4°C and then resuspended in 800 µl of EasyPure Total RNA reagent (Bioman). After adding 20 µg of glycogen (Roche), RNA was purified according to the manufacturer’s instructions.

#### P. marinus – ammocoetes

Total RNA was extracted using the RNeasy Mini Kit (Qiagen), following the manufacturer’s protocol. An on-column DNase treatment (Qiagen) was performed to eliminate residual genomic DNA.

### Expression analysis

#### S. purpuratus (larva)

Extracted RNA was used to construct libraries that were sequenced on an Illumina HiSeq2000 to generate 150 bp paired-end reads. Raw sequencing data has been deposited in NCBI (XXXAccNo_TBDXXX).

#### S. purpuratus (adult)

Using Superscript IV reverse transcriptase (Invitrogen) approximately 100 ng of total RNA were converted to cDNA according to the manufacturer’s guidelines. The cDNA products were diluted 1:1 with water before being used as templates in the subsequent PCR reactions.

The quantification of transcript levels was performed using an ABI 7500 FAST realtime PCR machine. Ten microliter reactions containing 5 µl Fast SYBR Green Master Mix (ABI, 4385612) and 1 µl cDNA were set up in triplicate for each sample for each time point using the qPCR primer pairs listed in Suppl. Table 3. The amounts of *SpurIL-1* and *SpurIL-17-9* transcripts were normalized to that of 18S in each sample.

#### *P. marinus* (ammocoetes)

First-strand cDNA synthesis was carried out using the SuperScript™ IV First-Strand Synthesis System (ThermoFisher) with 1 µg of total RNA. Quantitative real-time PCR (qPCR) was conducted using SYBR Green chemistry on an Applied Biosystems QuantStudio 7 Flex Real-Time PCR System with the primer pairs listed in Suppl. Table 3. Each sample was run in triplicate to ensure technical reproducibility. Gene expression levels were normalized to that of the reference gene β-actin (LOC116951639).

### Expression analysis using HCR FISH

#### S. purpuratus (larva)

Larvae were fixed in 4% PFA for 1 hr at RT, washed three times in PBST (PBS, 0.1% Tween-20), dehydrated in ethanol and stored in 70% ethanol at −20°C prior to HCR. All subsequent steps were conducted according to the manufacturer’s protocol.

#### S. purpuratus (adult)

HCR RNA-FISH (Molecular instruments) was done according to the manufacturers protocol with minor modifications. Briefly, coelomocytes were collected from apparently healthy adult *S. purpuratus* (as described in the RNA preparation section), diluted to 150,000 cells/ml with CMFSW-EH, and aliquots of 2 ml cell suspension were plated onto pre-rinsed 22 mm diameter coverslips in a matching container. After letting the cells settle down for 30 min, they were first fixed with 4% paraformaldehyde in PBS at room temperature for 10 min, followed by three washes with PBS, and then fixed a second time with 70% ethanol at 4°C for 30 min. The coverslips were then washed with 2× SSC (0.3 M NaCl, 0.30 M sodium citrate, pH 7.0) and pre-hybridized with 100 µl HCR hybridization buffer (Molecular Instruments) at 37°C for 30 min, followed by the addition of 0.4 µl HCR probe and hybridization at 37°C for 18-20 h. Subsequently, the hybridization mix was removed, and the samples were washed three times with HCR wash buffer for 5 min, and three times with 5× SSCT (5× SSC [0.75M NaCl, 0.075M sodium citrate] containing 0.1% Tween 20) for 5 min. After a 30 min incubation with HCR amplification buffer, HCR amplifiers (B1-488, 1 µl of each hairpin in 50 µl) were prepared, denatured, cooled, added to the samples, and incubated at room temperature for 18-20 h. Then the hairpin mix was removed, and the samples were again washed three times with 5× SSCT. Finally, the nuclei were stained with 330 nM 2 -(4-Amidinophenyl)-6-indolecarbamidine (DAPI) in PBS for 5 min, and the coverslips were washed again three times with PBS for 5 min.

### Analysis of RNA-Seq datasets

#### S. purpuratus (larva)

To analyze the sea urchin larval immune response, the read quality of the raw data (see above) was assessed using FastQC^10^ and reads were trimmed using TrimGalore, v0.6.6 (github.com/FelixKrueger/TrimGalore). Reads were mapped onto the *S. purpuratus* genome sequence (v5.0) obtained from Echinobase^14^ using Hi-Sat2 v2.0.5^11^; transcript counts were calculated using StringTie^12^. Counts were normalized across samples by calculating RPKM values based on total numbers of mapped reads and transcript length using custom scripts. Differential expression analysis was performed using DESeq2^13^.

#### S. purpuratus (adult tissues)

The raw Illumina reads from adult gut (SRR531955), ovary (SRR531958), radial nerve (SRR532046), testes (SRR532121), axial gland (SRR531951) and coelomocytes (SRR531953) reads^16^ were mapped onto the Spur5.0 genome (GCF_000002235.5) using TopHat2^17^ with default parameters and the expression levels of individual genes were calculated using cuffdiff^18^ with default parameters and the gene annotation file available at GenBank (GCF_000002235.5_Spur_5.0_genomic.gff).

#### N. vectensis

To investigate the possibility that *IL-1anc* transcripts are regulated in the cnidarian immune response, the raw sequencing files were obtained from NCBI (PRJNA734503; ^15^). Reads were mapped against the *N. vectensis* genome ([jaNemVect1.1](https://www.ncbi.nlm.nih.gov/datasets/genome/GCF_932526225.1/); GCA_932526225.1) and analyzed using the pipeline described for the *S. purpuratus* larva datasets.

### Cloning of the IL-1 transcripts

#### S. purpuratus

The ORFs of each *SpurIL-1* gene were amplified from coelomocyte cDNA using Phusion polymerase and gene specific primers (Suppl. Table 4). The products were gel-purifiedusing the EasyPure PCR/Gel extraction kit (Bioman), cloned into the pJET1.2 vector using the CloneJET PCR Cloning Kit (Thermo Scientific). Positive clones were identified by restriction digest and the identity of the inserts was confirmed by Sanger sequencing.

#### P. marinus

The cDNAs encoding PmarIL-1.1 (XP_032830942.1) and PmarIL-1.2 (XP_032830949.1) codon-optimized for expression in human cell lines were synthesized by a commercial service (Twist Bioscience). The cDNA encoding PmarIL-1.3 (XP_032830829.1) codon-optimized for expression in human cell lines and with a C-terminal 3×FLAG tag was synthesized and cloned into the pTwistCMV mammalian expression vector (Twist Bioscience).

### Generation of IL-1 expression vectors

The pCMV_HsIL1b-FLAG-IRES-GFP expression vector for the human pro-IL1β (NP_000567.1) was a gift from Dr. Ming-Zong Lai (Academia Sinica, Taipei, Taiwan). To generate expression vectors for the FLAG-tagged *S. purpuratus* IL-1s we amplified the complete ORFs of each transcript using Phusion polymerase with respective primer pairs (Supplementary Table 2) such that the products harbored SpeI and SalI (SpurIL-1.1, SpurIL-1.2), BamHI and SalI (SpurIL-1.3, SpurIL-1.5), BamHI and XhoI (SpurIL-1.4), or SpeI and XhoI (PmarIL-1.1, PmarIL-1.2) restriction sites at their 5’ and 3’ ends, respectively. These fragments were then digested with these enzymes and ligated into pCMV-FLAG-IRES-GFP that was digested with SpeI+XhoI or BamHI+XhoI, respectively. Positive clones were confirmed by Sanger sequencing.

### Site-directed mutagenesis of IL-1 expression vectors

To change individual amino acids within the SpurIL-1.3 protein encoded by the pCMV_SpurIL-1.3-FLAG-IRES-GFP expression vector the Quikchange approach (Stratagene) was used. Complementary top and bottom strand oligonucleotides harboring the respective mutations(Suppl. Table 5) were used as primers in a 50 µl PCR reaction (98°C, 2 min; 16×[98°C, 20 s; 55°C, 20 s; 72°C; 3.5 min]; 72°C, 5 min) using Phusion polymerase and the respective wildtype expression vectors as the template. The entire PCR reactions were subsequently digested with DpnI at 37°C for 30 min, and aliquots of this digest were transformed into competent DH10B *E. coli*. The entire inserts of individual clones were sequenced to confirm that the desired mutation is present, and that no additional off-target mutations were introduced.

### Generation of Caspase-1 expression vectors

Human Caspase1 (HsCASP1, NP_001214.1) cDNA was a gift from Dr. Ming -Zong Lai (Academia Sinica, Taipei, Taiwan) and the *P. marinus* Caspase1 (PmCasp1, XP_032814408.1) cDNA was synthesized as codon-optimized for expression in human cell lines by a commercial service (Twist Bioscience). The complete ORFs were amplified by PCR with gene-specific primers (Supplementary Table 2) using Phusion polymerase and cloned into the pCDNA3.1-V5/His TOPO mammalian expression vector (Invitrogen). The plasmid for expressing the HsCASP1 C285A catalytic mutant was generatedby site-directed mutagenesis (see previous section) using the primers indicated in Supplementary Table 3.

### Tissue culture

Human 293T cells were maintained in Dulbecco’s Modified Eagle Medium (DMEM, high glucose, Gibco 11965-092) containing 10% fetal bovine serum (Cytiva, SH30396.13), 1× Penicillin (100 units/ml)/Streptomycin (0.1mg/ml) (Capricorn Scientific), at 37°C in a humidified 5% CO2 atmosphere.

### Transfections

Aliquots of 250,000 293T cells were seeded into individual wells of a 6-well plate containing 2 ml of growth medium. After incubation for 16–24 hours at 37°C in a 5% CO₂, the cells were transfected using Lipofectamine™ 3000 (Invitrogen) following the manufacturer’s protocol. Briefly, the transfection mixture comprised 3.75 µl Lipofectamine™ 3000, 1 µl P3000 reagent, and 1 µg plasmid DNA diluted in Opti-MEM™ I Reduced Serum Medium (Gibco 31985-062). After adding this mix to the culture, cells were incubated for an additional 16 – 24 hours under the same conditions.

### Western blot

Post-incubation, the growth medium was aspirated, the cells were washed once with ice-cold 1× PBS, and 200 µl ice-cold RIPA buffer (150 mM NaCl, 1% Triton X-100, 0.5% sodium deoxycholate, and 0.1% SDS in 50 mM Tris-HCl pH 8.0) including protease inhibitors (1× Aprotinin, 1× Leupeptin, 1 mg/ml Pepstatin A, and 1 mM PMSF) were added to each well. After incubating on ice with gentle shaking for 10 min, the lysates were transferred into 1.5 ml microcentrifuge tubes and the insoluble debris was pelleted at 12,000×g at 4°C for 1 minute. The supernatants were mixed with an equal volume of 2× SDS gelloading buffer (4% SDS, 20% glycerol, 200mM DTT, 0.01% bromophenol blue and 0.125 M Tris HCl pH=6.8) and the protein were denatured at 95°C for 10 min. Aliquots were loaded onto 12% SDS/PAGE gels, and separated at 50 V for 10 min, 80 V for 35 minutes, and finally at 120 V for 50 min. Afterwards the proteins were transferred to PDVF membranes using semidry transfer at 30V for 40 min.

To visualize the protein bands, the PVDF membrane was rinsed once with 1× TBST (Tris-buffered saline containing 0.1% Tween 20 (TBST) and incubated in blocking buffer (5% non-fat dry milk in 1× TBST) at room temperature for 1 h. The membrane was then incubated overnight at 4°C with 5 ml of primary antibody solution (mouse Anti-V5 [ab27671, Abcam], 1:10,000 dilution, mouse Anti-FLAG M2 [F3165, Merck], 1:10000 dilution, or mouse anti-actin C4 [MAB1501, Sigma] 1:20,000 in blocking buffer) and washed three times with 1× TBST for 5 min. The membrane was then incubated with 5 ml of secondary antibody (Merck, AP124P, goat anti-mouse HRP-conjugated, 1:10,000 dilution in 5% milk/TBST) for 1 hour at room temperature followed again by three washes with 1× TBST. Finally, the membrane was incubated with ECL reagents (T-Pro Biotech) according to the manufacturer’s instruction and the chemiluminescene images were captured using an Amersham IQ800 imaging system (Cytiva).

### Immunofluorescence

One day after transfection, the growth medium was removed, the cells were fixed with 4% paraformaldehyde in PBS for 10 min and permeabilized by incubation with 0.25% Triton X-100 in PBS for 10 min. After three washes with PBS for 5 min, the coverslips were blocked with 3% bovine serum albumin (BSA) in PBS for 30 min, and then incubated with the primary antibody (mouse Anti-FLAG M2 [F3165, Merck], 1:1000 dilution) in PBS containing 3% BSA for 30 min. After washing three times with PBS for 5 min, the coverslips were incubated in the dark with an Alexa 594-coupled goat α-mouse IgG antiserum (1:1000 dilution, [115-585-003, Jackson lab]) in PBS containing 2% newborn goat serum (NGS) for 1 h. After three washes with PBS for 5 min, the nuclei were stained with 330 nM DAPI in PBS for 5 min. Finally, the coverslips were washed again three times with PBS for 5 min.

### Microscopy and image acquisition

#### Fixed *S. purpuratus* coelomocytes and 293T cells

Coverslips containing fixed samples were mounted onto glass slides using 12 μl of 1,4-Diazobicyclo-(2,2,2-octane) (DABCO) mounting medium(2.5% solution in 70% glycerol with 10 mM Tris pH=7.5) and sealed with nail polish. Phase-contrast and fluorescence light images were taken using a Zeiss Axioimager.Z2 microscope using a cooled CCD camera. The raw image files were processed using the Zeiss Zen software.

#### Fixed S. purpuratus larvae

To image, fixed larvae were wet-mounted onto glass slides using coverslips suspended by clay feet. DIC and fluorescent images were captured using a Zeiss Axioimager.

### Synteny analysis

Macrosynteny analysis was performed as described^19^. Unique orthologs among species were identified using OrthoFinder^20^; the resulting data were used as input to construct ribbon diagrams for selected species. Figures were generated using MacrosyntR^21^. Significant associations were computed using Fisher’s exact test.

### Statistical analysis

The statistical analysis of the qPCR datasets was done using GraphPad prism.

## References

1. Beck, G., O’Brien, R. F., Habicht, G. S., Stillman, D. L., Cooper, E. L., and Raftos, D. A. (1993). Invertebrate cytokines. III: Invertebrate interleukin-1-like molecules stimulate phagocytosis by tunicate and echinoderm cells. Cell Immunol 146, 284–299. doi: S0008-8749(83)71027-0 [pii] 10.1006/cimm.1993.1027

2. Bittrich, S., Segura, J., Duarte, J. M., Burley, S. K., and Rose, Y. (2024). RCSB protein Data Bank: exploring protein 3D similarities via comprehensive structural alignments. Bioinformatics 40, btae370. doi: 10.1093/bioinformatics/btae370

3. Boraschi, D. (2022). What Is IL-1 for? The Functions of Interleukin-1 Across Evolution. Front Immunol 13, 872155. doi: 10.3389/fimmu.2022.872155

4. Boraschi, D., Italiani, P., Weil, S., and Martin, M. U. (2018). The family of the interleukin-1 receptors. Immunol Rev 281, 197–232. doi: 10.1111/imr.12606

5. Boulay, J.-L., Du Pasquier, L., and Cooper, M. D. (2022). Cytokine Receptor Diversity in the Lamprey Predicts the Minimal Essential Cytokine Networks of Vertebrates. The Journal of Immunology 209, 1013–1020. doi: 10.4049/jimmunol.2200274

6. Brodsky, I. E., and Monack, D. (2009). NLR-mediated control of inflammasome assembly in the host response against bacterial pathogens. Seminars in immunology. doi: S1044-5323(09)00051-7 [pii] 10.1016/j.smim.2009.05.007

7. Buckley, K. M., Ho, E. C. H., Hibino, T., Schrankel, C. S., Schuh, N. W., Wang, G., et al. (2017). IL17 factors are early regulators in the gut epithelium during inflammatory response to Vibrio in the sea urchin larva. eLife 6. doi: 10.7554/eLife.23481

8. Buckley, K. M., and Rast, J. P. (2015). Diversity of animal immune receptors and the origins of recognition complexity in the deuterostomes. Developmental and Comparative Immunology 49, 179–189. doi: 10.1016/j.dci.2014.10.013

9. Chen, S., Li, S., Chen, H., Gong, Y., Yang, D., Zhang, Y., et al. (2023). Caspase-mediated LPS sensing and pyroptosis signaling in Hydra. Sci Adv 9, eadh4054. doi: 10.1126/sciadv.adh4054

10. Corradi, A., Bajetto, A., Cozzolino, F., and Rubartelli, A. (1993). Production and secretion of interleukin 1 receptor antagonist in monocytes and keratinocytes. Cytotechnology 11 Suppl 1, S50–52.

11. Degnan, S. M. (2015). The surprisingly complex immune gene repertoire of a simple sponge, exemplified by the NLR genes: A capacity for specificity? Developmental & Comparative Immunology 48, 269–274. doi: 10.1016/j.dci.2014.07.012

12. Di Paolo, N. C., and Shayakhmetov, D. M. (2016). Interleukin 1α and the inflammatory process. Nat Immunol 17, 906–913. doi: 10.1038/ni.3503

13. Dijkstra, J. M. (2014). TH2 and Treg candidate genes in elephant shark. Nature 511, E7–9. doi: 10.1038/nature13446

14. Dinarello, C. A. (2018). Overview of the IL -1 family in innate inflammation and acquired immunity. Immunological Reviews 281, 8–27. doi: 10.1111/imr.12621

15. Eggestøl, H. Ø., Lunde, H. S., Knutsen, T. M., and Haugland, G. T. (2020). Interleukin-1 Ligands and Receptors in Lumpfish (Cyclopterus lumpus L.): Molecular Characterization, Phylogeny, Gene Expression, and Transcriptome Analyses. Front. Immunol. 11, 502. doi: 10.3389/fimmu.2020.00502

16. Evavold, C. L., Ruan, J., Tan, Y., Xia, S., Wu, H., and Kagan, J. C. (2018). The Pore-Forming Protein Gasdermin D Regulates Interleukin-1 Secretion from Living Macrophages. Immunity 48, 35–44.e6. doi: 10.1016/j.immuni.2017.11.013

17. Faham, S., Linhardt, R. J., and Rees, D. C. (1998). Diversity does make a difference: fibroblast growth factor-heparin interactions. Curr Opin Struct Biol 8, 578–586. doi: 10.1016/s0959-440x(98)80147-4

18. Ghosh, J., Buckley, K. M., Nair, S. V., Raftos, D. A., Miller, C. A., Majeske, A. J., et al. (2010). Sp185/333: A novel family of genes and proteins involved in the purple sea urchin immune response. Developmental and comparative immunology 34, 235–45.

19. Heilig, R., Dick, M. S., Sborgi, L., Meunier, E., Hiller, S., and Broz, P. (2018). The Gasdermin-D pore acts as a conduit for IL-1β secretion in mice. Eur J Immunol 48, 584–592. doi: 10.1002/eji.201747404

20. Hibino, T., Loza-Coll, M., Messier, C., Majeske, A. J., Cohen, A. H., Terwilliger, D. P., et al. (2006). The immune gene repertoire encoded in the purple sea urchin genome. Developmental Biology 300. doi: 10.1016/j.ydbio.2006.08.065

21. Ho, E. C. H., Buckley, K. M., Schrankel, C. S., Schuh, N. W., Hibino, T., Solek, C. M., et al. (2016). Perturbation of gut bacteria induces a coordinated cellular immune response in the purple sea urchin larva. Immunology and Cell Biology 94, 861–874. doi: 10.1038/icb.2016.51

22. Jiang, S., Zhou, Z., Sun, Y., Zhang, T., and Sun, L. (2020). Coral gasdermin triggers pyroptosis. Sci. Immunol. 5, eabd2591. doi: 10.1126/sciimmunol.abd2591

23. Jumper, J., Evans, R., Pritzel, A., Green, T., Figurnov, M., Ronneberger, O., et al. (2021). Highly accurate protein structure prediction with AlphaFold. Nature 596, 583–589. doi: 10.1038/s41586-021-03819-2

24. Kaplanski, G., Farnarier, C., Kaplanski, S., Porat, R., Shapiro, L., Bongrand, P., et al. (1994). Interleukin-1 induces interleukin-8 secretion from endothelial cells by a juxtacrine mechanism. Blood 84, 4242–4248.

25. Kim, B., Lee, Y., Kim, E., Kwak, A., Ryoo, S., Bae, S. H., et al. (2013). The Interleukin-1α Precursor is Biologically Active and is Likely a Key Alarmin in the IL-1 Family of Cytokines. Front Immunol 4, 391. doi: 10.3389/fimmu.2013.00391

26. Krumm, B., Meng, X., Li, Y., Xiang, Y., and Deng, J. (2008). Structural basis for antagonism of human interleukin 18 by poxvirus interleukin 18-binding protein. Proc Natl Acad Sci U S A 105, 20711–20715. doi: 10.1073/pnas.0809086106

27. Krumm, B., Xiang, Y., and Deng, J. (2014). Structural biology of the IL-1 superfamily: Key cytokines in the regulation of immune and inflammatory responses. Protein Science 23, 526–538. doi: 10.1002/pro.2441

28. Kurt-Jones, E. A., Beller, D. I., Mizel, S. B., and Unanue, E. R. (1985). Identification of a membrane-associated interleukin 1 in macrophages. Proc Natl Acad Sci U S A 82, 1204–1208. doi: 10.1073/pnas.82.4.1204

29. Li, J.-H., Shao, J.-Z., Xiang, L.-X., and Wen, Y. (2007). Cloning, characterization and expression analysis of pufferfish interleukin-4 cDNA: the first evidence of Th2-type cytokine in fish. Mol Immunol 44, 2078–2086. doi: 10.1016/j.molimm.2006.09.010

30. Lin, Z., Akin, H., Rao, R., Hie, B., Zhu, Z., Lu, W., et al. (2023). Evolutionary-scale prediction of atomic-level protein structure with a language model. Science 379, 1123–1130. doi: 10.1126/science.ade2574

31. Luheshi, N. M., Rothwell, N. J., and Brough, D. (2009). The dynamics and mechanisms of interleukin-1alpha and beta nuclear import. Traffic 10, 16–25. doi: 10.1111/j.1600-0854.2008.00840.x

32. Margolis, S. R., Dietzen, P. A., Hayes, B. M., Wilson, S. C., Remick, B. C., Chou, S., et al. (2021). The cyclic dinucleotide 2′3′-cGAMP induces a broad antibacterial and antiviral response in the sea anemone *Nematostella vectensis*. Proc. Natl. Acad. Sci. U.S.A. 118, e2109022118. doi: 10.1073/pnas.2109022118

33. Mariotti, F. R., Supino, D., Landolina, N., Garlanda, C., Mantovani, A., Moretta, L., et al. (2023). IL-1R8: A molecular brake of anti-tumor and anti-viral activity of NK cells and ILC. Seminars in Immunology 66, 101712. doi: 10.1016/j.smim.2023.101712

34. Marlétaz, F., Timoshevskaya, N., Timoshevskiy, V. A., Parey, E., Simakov, O., Gavriouchkina, D., et al. (2024). The hagfish genome and the evolution of vertebrates. Nature 627, 811–820. doi: 10.1038/s41586-024-07070-3

35. Matsushima, K., Taguchi, M., Kovacs, E. J., Young, H. A., and Oppenheim, J. J. (1986). Intracellular localization of human monocyte associated interleukin 1 (IL 1) activity and release of biologically active IL 1 from monocytes by trypsin and plasmin. J Immunol 136, 2883–2891.

36. Meng, E. C., Pettersen, E. F., Couch, G. S., Huang, C. C., and Ferrin, T. E. (2006). Tools for integrated sequence-structure analysis with UCSF Chimera. BMC Bioinformatics 7, 339. doi: 10.1186/1471-2105-7-339

37. Oeckinghaus, A., and Ghosh, S. (2009). The NF-kappaB family of transcription factors and its regulation. Cold Spring Harbor perspectives in biology 1, a000034. doi: 10.1101/cshperspect.a000034

38. Ottaviani, E., Franchini, A., and Franceschi, C. (1993). Presence of Several Cytokine-like Molecules in Molluscan Hemocytes. Biochemical and Biophysical Research Communications 195, 984–988. doi: 10.1006/bbrc.1993.2141

39. Qin, K., Jiang, S., Xu, H., Yuan, Z., and Sun, L. (2024). Pyroptotic gasdermin exists in Mollusca and is vital to eliminating bacterial infection. Cell Rep 43, 114644. doi: 10.1016/j.celrep.2024.114644

40. Rast, J. P., and Messier-Solek, C. (2008). Marine invertebrate genome sequences and our evolving understanding of animal immunity. Biological Bulletin 214, 274–283. doi: 214/3/274 [pii]

41. Rivers-Auty, J., Daniels, M. J. D., Colliver, I., Robertson, D. L., and Brough, D. (2018). Redefining the ancestral origins of the interleukin-1 superfamily. Nat Commun 9, 1156. doi: 10.1038/s41467-018-03362-1

42. Roberts, S., Gueguen, Y., de Lorgeril, J., and Goetz, F. (2008). Rapid accumulation of an interleukin 17 homolog transcript in Crassostrea gigas hemocytes following bacterial exposure. Developmental and comparative immunology 32, 1099–1104. doi: S0145-305X(08)00045-1 [pii] 10.1016/j.dci.2008.02.006

43. Robertson, A. J., Croce, J. C., Carbonneau, S., Voronina, E., Miranda, E., McClay, D. R., et al. (2006). The genomic underpinnings of apoptosis in Strongylocentrotus purpuratus. Developmental Biology 300, 321–334.

44. Rubartelli, A., Cozzolino, F., Talio, M., and Sitia, R. (1990). A novel secretory pathway for interleukin-1 beta, a protein lacking a signal sequence. EMBO J 9, 1503–1510. doi: 10.1002/j.1460-2075.1990.tb08268.x

45. Schreuder, H., Tardif, C., Trump-Kallmeyer, S., Soffientini, A., Sarubbi, E., Akeson, A., et al. (1997). A new cytokine-receptor binding mode revealed by the crystal structure of the IL-1 receptor with an antagonist. Nature 386, 194–200. doi: 10.1038/386194a0

46. Schroder, K., and Tschopp, J. (2010). The inflammasomes. Cell 140, 821–32. doi: 10.1016/j.cell.2010.01.040

47. Smith, J. J., Timoshevskaya, N., Ye, C., Holt, C., Keinath, M. C., Parker, H. J., et al. (2018). The sea lamprey germline genome provides insights into programmed genome rearrangement and vertebrate evolution. Nat Genet 50, 270–277. doi: 10.1038/s41588-017-0036-1

48. Tu, Q., Cameron, R. A., Worley, K. C., Gibbs, R. a, and Davidson, E. H. (2012). Gene structure in the sea urchin Strongylocentrotus purpuratus based on transcriptome analysis. Genome research 22, 2079–87. doi: 10.1101/gr.139170.112

49. Van Kempen, M., Kim, S. S., Tumescheit, C., Mirdita, M., Lee, J., Gilchrist, C. L. M., et al. (2023). Fast and accurate protein structure search with Foldseek. Nat Biotechnol. doi: 10.1038/s41587-023-01773-0

50. Vinkler, M., Fiddaman, S. R., Těšický, M., O’Connor, E. A., Savage, A. E., Lenz, T. L., et al. (2023). Understanding the evolution of immune genes in jawed vertebrates. J Evol Biol 36, 847–873. doi: 10.1111/jeb.14181

51. Wang, W., Li, D., Luo, K., Chen, B., Hao, T., Li, X., et al. (2025). IL-1 Superfamily Across 400+ Species: Therapeutic Targets and Disease Implications. Biology 14, 561. doi: 10.3390/biology14050561

52. Wang, X., Wei, X., Lu, Y., Wang, Q., Fu, R., Wang, Y., et al. (2023). Characterization of GSDME in amphioxus provides insights into the functional evolution of GSDM-mediated pyroptosis. PLoS Biol 21, e3002062. doi: 10.1371/journal.pbio.3002062

53. Watthanasurorot, A., Söderhäll, K., Jiravanichpaisal, P., and Söderhäll, I. (2011). An ancient cytokine, astakine, mediates circadian regulation of invertebrate hematopoiesis. Cell Mol Life Sci 68, 315–323. doi: 10.1007/s00018-010-0458-8

54. Wu, S.-Z., Huang, X.-D., Li, Q., and He, M.-X. (2013). Interleukin-17 in pearl oyster (Pinctada fucata): molecular cloning and functional characterization. Fish & shellfish immunology 34, 1050–6. doi: 10.1016/j.fsi.2013.01.005

55. Xiang, Y., and Moss, B. (1999). IL-18 binding and inhibition of interferon gamma induction by human poxvirus-encoded proteins. Proc Natl Acad Sci U S A 96, 11537–11542. doi: 10.1073/pnas.96.20.11537

56. Xu, L., Yuan, S., Li, J., Ruan, J., Huang, S., Yang, M., et al. (2011). The conservation and uniqueness of the caspase family in the basal chordate, amphioxus. BMC Biol 9, 60. doi: 10.1186/1741-7007-9-60

57. Yuen, B., Bayes, J. M., and Degnan, S. M. (2014). The characterization of sponge NLRs provides insight into the origin and evolution of this innate immune gene family in animals. Molecular biology and evolution 31, 106–20. doi: 10.1093/molbev/mst174

58. Zandawala, M., and Gera, J. (2024). Leptin- and cytokine-like unpaired signaling in Drosophila. Mol Cell Endocrinol 584, 112165. doi: 10.1016/j.mce.2024.112165

59. Zhang, Q., Zmasek, C. M., and Godzik, A. (2010). Domain architecture evolution of pattern-recognition receptors. Immunogenetics 62, 263–272.

60. Zhao, T., and Schranz, M. E. (2019). Network-based microsynteny analysis identifies major differences and genomic outliers in mammalian and angiosperm genomes. Proc Natl Acad Sci U S A 116, 2165–2174. doi: 10.1073/pnas.1801757116

61. Zhong, M., Yan, H., and Li, Y. (2017). Flagellin: a unique microbe-associated molecular pattern and a multi-faceted immunomodulator. Cell Mol Immunol 14, 862–864. doi: 10.1038/cmi.2017.78

62. Zhu, X., Zhang, Z., Ren, J., Jia, L., Ding, S., Pu, J., et al. (2020). Molecular Characterization and Chemotactic Function of CXCL8 in Northeast Chinese Lamprey (Lethenteron morii). Front Immunol 11, 1738. doi: 10.3389/fimmu.2020.01738

## References

1. Eddy, S. R. HMMER User ‘ s Guide. 0–93 (2010).

2. Kim, G. et al. Easy and accurate protein structure prediction using ColabFold. Nat. Protoc. 20, 620–642 (2025).

3. Lin, Z. et al. Evolutionary-scale prediction of atomic-level protein structure with a language model. Science 379, 1123–1130 (2023).

4. Pettersen, E. F. et al. UCSF Chimera--a visualization system for exploratory research and analysis. J. Comput. Chem. 25, 1605–1612 (2004).

5. Bittrich, S., Segura, J., Duarte, J. M., Burley, S. K. & Rose, Y. RCSB protein Data Bank: exploring protein 3D similarities via comprehensive structural alignments. Bioinforma. Oxf. Engl. 40, btae370 (2024).

6. Zhang, Y. & Skolnick, J. TM-align: a protein structure alignment algorithm based on the TM-score. Nucleic Acids Res. 33, 2302–2309 (2005).

7. Hiller, K., Grote, A., Scheer, M., Münch, R. & Jahn, D. PrediSi: prediction of signal peptides and their cleavage positions. Nucleic Acids Res. 32, W375–379 (2004).

8. Ho, E. C. H. et al. Perturbation of gut bacteria induces a coordinated cellular immune response in the purple sea urchin larva. Immunol. Cell Biol. 94, 861–874 (2016).

9. Liu, M., Liao, W., Buckley, K. M., Yang, S. Y. & Fugmann, S. D. AID/APOBEC-like cytidine deaminases are ancient innate immune mediators in invertebrates. Nat. Commun. 9, 1–11 (2018).

10. Andrews, S. FastQC. (2010).

11. Kim, D., Paggi, J. M., Park, C., Bennett, C. & Salzberg, S. L. Graph-based genome alignment and genotyping with HISAT2 and HISAT-genotype. Nat. Biotechnol. 37, 907–915 (2019).

12. Pertea, M., Kim, D., Pertea, G. M., Leek, J. T. & Salzberg, S. L. Transcript-level expression analysis of RNA-seq experiments with HISAT, StringTie and Ballgown. Nat. Protoc. 11, 1650–67 (2016).

13. Love, M. I., Anders, S. & Huber, W. Differential Analysis of Count Data - the DESeq2 Package. (2014). doi:10.1101/002832.

14. Telmer, C. A. et al. Echinobase: a resource to support the echinoderm research community. GENETICS 227, iyae002 (2024).

15. Margolis, S. R., et al. The cyclic dinucleotide 2′3′-cGAMP induces a broad antibacterial and antiviral response in the sea anemone Nematostella vectensis. Proc. Natl. Acad. Sci. 118, e2109022118 (2021).

16. Tu, Q., Cameron, R. A., Worley, K. C., Gibbs, R. a & Davidson, E. H. Gene structure in the sea urchin Strongylocentrotus purpuratus based on transcriptome analysis. Genome Res. 22, 2079–87 (2012).

17. Trapnell, C., Pachter, L. & Salzberg, S. L. TopHat: discovering splice junctions with RNA-Seq. Bioinformatics 25, 1105–11 (2009).

18. Trapnell, C. et al. Transcript assembly and quantification by RNA-Seq reveals unannotated transcripts and isoform switching during cell differentiation. Nat. Biotechnol. 28, 511–5 (2010).

19. Simakov, O. et al. Deeply conserved synteny resolves early events in vertebrate evolution. Nat. Ecol. Evol. 4, 820–830 (2020).

20. Emms, D. M. & Kelly, S. OrthoFinder: phylogenetic orthology inference for comparative genomics. Genome Biol. 20, 238 (2019).

21. El Hilali, S. & Copley, R. R. macrosyntR : Drawing automatically ordered Oxford Grids from standard genomic files in R. Preprint at 10.1101/2023.01.26.525673 (2023).

