## Supplementary material for "An ancient evolutionary origin for IL-1 cytokines as mediators of immunity": Supplemenal Figures and Tables

Francisco Fontenla-Iglesias et al.

**Supplementary Figure S1. Alignment of metazoan IL-1 trefoil domain sequences based on pairwise structure alignment of the human IL-1β crystal structure (1ITB) and computed structure models of representative metazoan IL-1 proteins.** Although their primary sequence varies considerably, twelve β-strands (yellow highlight) are identified in each structure. Some small stretches of α-helix are also found in all or many structure models (designated A, blue highlight). The often conserved "FES" motif in β-strand 9 is shown in bold. All human sequences are from the designated crystal structures (except for IL-33, which is derived from a computational structure to account for unresolved residues in the available crystal structure). An unresolved lysine residue in the Human IL-37 structure (5HN1.A) is shown in green. Red amino acid characters indicate that although Chimera did not call a β-strand, the peptide backbone closely overlays the human IL1β β-strand 12 in this region. Human FGF-1 (1RG8) is shown for comparison.

**Supplementary Figure S2. Computationally folded structure models for complete chains of three representative IL-1anc proteins.** (**A**) *Danio rerio* (zebrafish) DrerIL-1anc.1 (NP_001277347.1); (**B**) *Petromyzon marinus* PmarIL-1.2; (**C**) *Strongylocentrotus purpuratus* SpurIL-1.2; (**D**) *Nematostella vectensis* NvecIL-1. PDZ domains are shown in blue, and the C-terminal IL-1 trefoil domains in green. Linking regions between the domains are shown in orange. Non-domain N-terminal and C terminal portions are shown in white.

**Supplementary Figure S3. Secretion of ectopically expressed IL-1s from mammalian cells.** (**A**) Signal peptide prediction using PrediSi indicated presence of an unusually long 57 amino acid signal peptide in the agnathan PmarIL-1.1 and EataIL-1.1, but not for any other of the IL-1 proteins from *S. purpuratus* and other invertebrates. (**B+C**) Human 293T cells were transfected with expression vectors for the indicated FLAG-tagged *P. marinus* and *S. purpuratus* IL-1s with human IL-1β (HsapIL-1β) serving as a control. Total cell lysates and concentrated culture supernatants were resolved by SDS/PAGE and the proteins were visualized by Western blot using a monoclonal anti-FLAG antibody. Anti-actin antibodies were used to show comparable loading of protein amounts for the cell lysates. Note that low levels of PmarIL-1.3 and HsapIL-1β in the culture supernatant are likely due to low levels of spontaneous cell death in the culture. Theoretical sizes of the FLAG-tagged IL-1s are: PmarIL-1.1=33.0 kD (including signal peptide) or 26.9 kD (after cleavage of signal peptide); PmarIL-1.2=57.8 kD; PmarIL-1.3=47.9 kD; SpurIL-1.1=63.5 kD; SpurIL-1.2=55.2 kD; SpurIL-1.3=51.4 kD; mature SpurIL-1.3=30.6 kD; SpurIL-1.4=59.7 kD; SpurIL-1.5=55.1 kD; HsapIL-1β=34 kD, mature HsapIL-1β=20.6 kD.

**Supplementary Figure S4. Microsynteny of the ancestral IL-1 in selected vertebrate phyla.** A schematic representation of the chromosomal regions surrounding the three *IL-1anc* genes (IL-1.1 to IL-1.3, cyan boxes) of jawless vertebrates and the single *IL-1anc* in selected jawed vertebrates (IL-1, cyan boxes) and the phylogenetic relationship of these species is provided. The syntenic region from *Bufo bufo* (common toad) is shown as a representative for the majority of tetrapods that lost the *IL-1anc* gene. Selected genes encoding one-to-one orthologs present in multiple species are highlighted in color to emphasize the synteny between the genomic regions. The *ESD* gene that is linked to *IL-1anc* genes in many invertebrates (Fig. 3) is also linked to *IL-1anc* in two

*Chondrichthyes*. Note that the *Danio rerio* locus harbors two genes encoding IL-1 proteins, the complete IL-1anc.1 (IL-1,cyan) and a closely related IL-1anc.2 (IL-1x, cyan) that appears to have lost the PDZ domain.

**Supplementary Figure S5. Synteny of chromosomal regions harboring *IL-1anc* genes across metazoans.**
The *IL-1anc* genes (asterisks) are located on orthologous chromosomes in *N. vectensis* (chr9, starlet sea anemone), *P. maximus* (chr14, great scallop), *A. rubens* (chr13, great scallop), *Lepisosteus oculatus* (chr15, spotted gar), and *P. marinus* (chr55). The statistical significance of these associations ($p \leq 0.05$, Bonferroni-corrected one-sided Fisher's exact test) supports the presence of a conserved syntenic block including the *IL-1anc* genes that is maintained since the cnidarian–vertebrate last common ancestor. (**A–B**) Representative ribbon plots were generated using macrosyntR with (**A**) comparing *N. vectensis*, *P. maximus*, *A. rubens*, and *L. oculatus* and (**B**) comparing *N. vectensis*, *P. maximus*, *A. rubens*, and *P. marinus* to improve visual clarity. Each horizontal bar represents a chromosome and vertical connectors indicating individual orthologous genes, color-coded according to their corresponding ortholog in *N. vectensis*.

**Supplementary Figure S6. Crystal structure and computational models of the IL-1 trefoil domain from representative deuterostomes.** (**A**) Human IL-1β (1ITB); (**B**) Human IL-18 (3WO2); (**C**) Human IL-33 (4KC3); (**D**) Human IL-37 (5HN1); (**E**) *Petromyzon marinus* PmarIL-1.1; (**F**) *Petromyzon marinus* PmarIL-1.2; (**G**) *Petromyzon marinus* PmarIL-1.3; (**H**) *Eptatretus atami* EataIL-1.1; (**I**) *Eptatretus atami* EataIL-1.2; (**J**) *Eptatretus atami* EataIL-1.3; (**K**) *Branchiostoma floridae* BfloIL-1; (**L**) *Branchiostoma belcheri* BbelIL-1; (**M**) *Strongylocentrotus purpuratus* SpurIL-1.1; (**N**) *Strongylocentrotus purpuratus* SpurIL-1.2; (**O**) *Strongylocentrotus purpuratus* SpurIL-1.3; (**P**) *Strongylocentrotus purpuratus* SpurIL-1.4; (**Q**) *Strongylocentrotus purpuratus* SpurIL-1.5; (**R**) *Holothuria leucospilota* HleuIL-1.1; (**S**) *Holothuria leucospilota* HleuIL-1.2; (**T**) *Patiria miniata* PminIL-1.1; (**U**) *Patiria miniata* PminIL-1.2; (**V**) *Patiria miniata* PminIL-1.3; (**W**) *Patiria miniata* PminIL-1.4; (**X**) *Anneissia japonica* AjapIL-1.1; (**Y**) *Anneissia japonica* AjapIL-1.2; (**Z**) *Anneissia japonica* AjapIL-1.3; (**AA**) *Saccoglossus kowalevskii* SkowIL-1.2; (**BB**) *Ptychodera flava* PflaIL-1.1. Two images are shown for each model, with the second rotated 90° around the vertical axis. PDB IDs are given for the human crystal structure models. Other sequences are computational models derived with ESMFold.

**Supplementary Figure S7. Computational models of the IL-1 trefoil domain from representative protostomes and cnidarians.** (**A**) *Procambarus clarkii* (red swamp crayfish) PclaIL-1.1; (**B**) *Procambarus clarkii* PclaIL-1.2; (**C**) *Procambarus clarkii* PclaIL-1.3; (**D**) *Scylla paramamosain* (green mud crab) SparIL-1.1; (**E**) *Scylla paramamosain* SparIL-1.2; (**F**) *Penaeus vannamei* (Pacific white shrimp) PvanIL-1.1; (**G**) *Pecten maximus* (great scallop) PmaxIL-1.1; (**H**) *Pecten maximus* PmaxIL-1.2; (**I**) *Pecten maximus* PmaxIL-1.3; (**J**) *Crassostrea virginica* (American oyster) CvirIL-1.1; (**K**) *Crassostrea virginica* CvirIL-1.2; (**L**) *Lingula anatina* LanaIL-1.1; (**M**) *Lingula anatina* LanaIL-1.2; (**N**) *Exaiptasia diaphana* (brown anemone) EpalIL-1; (**O**) *Nematostella* *vectensis* (starlet sea anemone) NvecIL-1; (**P**) *Actinia tenebrosa* (waratah anemone) AtenIL-1. Two images are shown for each model, the second rotated 90° around the vertical axis.

**Supplementary Figure S8. Microsynteny of *IL-1anc* loci in additional invertebrate phyla.** A schematic representation of the chromosomal regions surrounding the *IL-1anc* genes (IL-1, cyan boxes) in selected (**A**) *Cephalochordata*, (**B**) *Lophotrochozoa*, and (**C**) *Crustacea* and the phylogenetic relationships are shown. Selected genes encoding one-to-one orthologs present in multiple species within each phylum are highlighted in color to emphasize the synteny between the genomic regions. Note that there is no discernible microsynteny within the *Lophotrochozoans*.

**Supplementary Figure S9. Intron positions in representative *IL-1* genes show conserved phasing in PDZ and IL-1 trefoil domains.** Phasing in **(A)** representative human *IL-1* family genes (for comparison), **(B)** the three *Petromyzon marinus IL-1* genes, **(C)** the five *Strongylocentrotus purpuratus IL-1* genes, and **(D)** in representative protostome and cnidarian *IL-1* genes. PDZ domains, when present, are shown as pink boxes and IL-1 trefoil domains are shown as blue boxes. The position of the conserved FES motif (or related sequence) in β-strand 12 is shown with an arrowhead. Note that intron and exons lengths are not drawn to scale.

**Supplementary Figure S10. Gene expression data from adult and larval infection experiments.** **(A)** The transcript levels of the *SpIL-17-9* genes were assessed in three adult *S. purpuratus* (same individuals as in Fig. 4) by qRT-PCR prior to infection with *V. diazotrophicus* (t=0 h) and six hours later (t=6 h). All values were normalized to the levels of 18S transcripts in each sample, and the data for each individual is shown separately. (**B-I**) Transcript levels of the indicated *S. purpuratus SpurIL-1* and control genes (*SpIL-17-1* and *185/333*) were quantified by RNAseq analysis in pooled larvae at the indicated time points prior to (t=0) and during an acute *V. diazotrophicus* infection. (**J-K**) Fluorescence *in situ* hybridization was conducted on fixed *S. purpuratus* larvae prior to (t=0 h) and after infection with *V. diazotrophicus* (t=6 h) using a pan-*SpurIL-1.1/2/3* HCR RNA probe labelled in green. The nuclei were stained with DAPI (blue), and images were collected using visual light differential interference contrast (DIC) and fluorescence microscopy. Representative images are shown separated by channel and merged (DIC+blue+green and blue+green).

**Supplementary Figure S11. Additional HCR-FISH data from adult *S. purpuratus* coelomocytes.** Adult *S. purpuratus* were infected with *V. diazotrophicus*, and the expression of the *SpurIL-1* genes in coelomocytes was assessed by HCR-FISH (green) on fixed cell samples collected prior to infection (t=0 h) and six hours later (t=6 h) using the indicates probe sets. While one probe set (SpurIL-1.1/2/3) cross-hybridizes with these three genes, those for SpurIL-1.4 and SpurIL-1.5 probes were gene-specific. The nuclei were stained with DAPI (blue), and images were collected using visual light phase contrast (ph) and fluorescence microscopy. Representative images are shown separated by channel and merged (ph+green and blue+green). The phagocytes were classified manually based on their morphology (colored arrows).

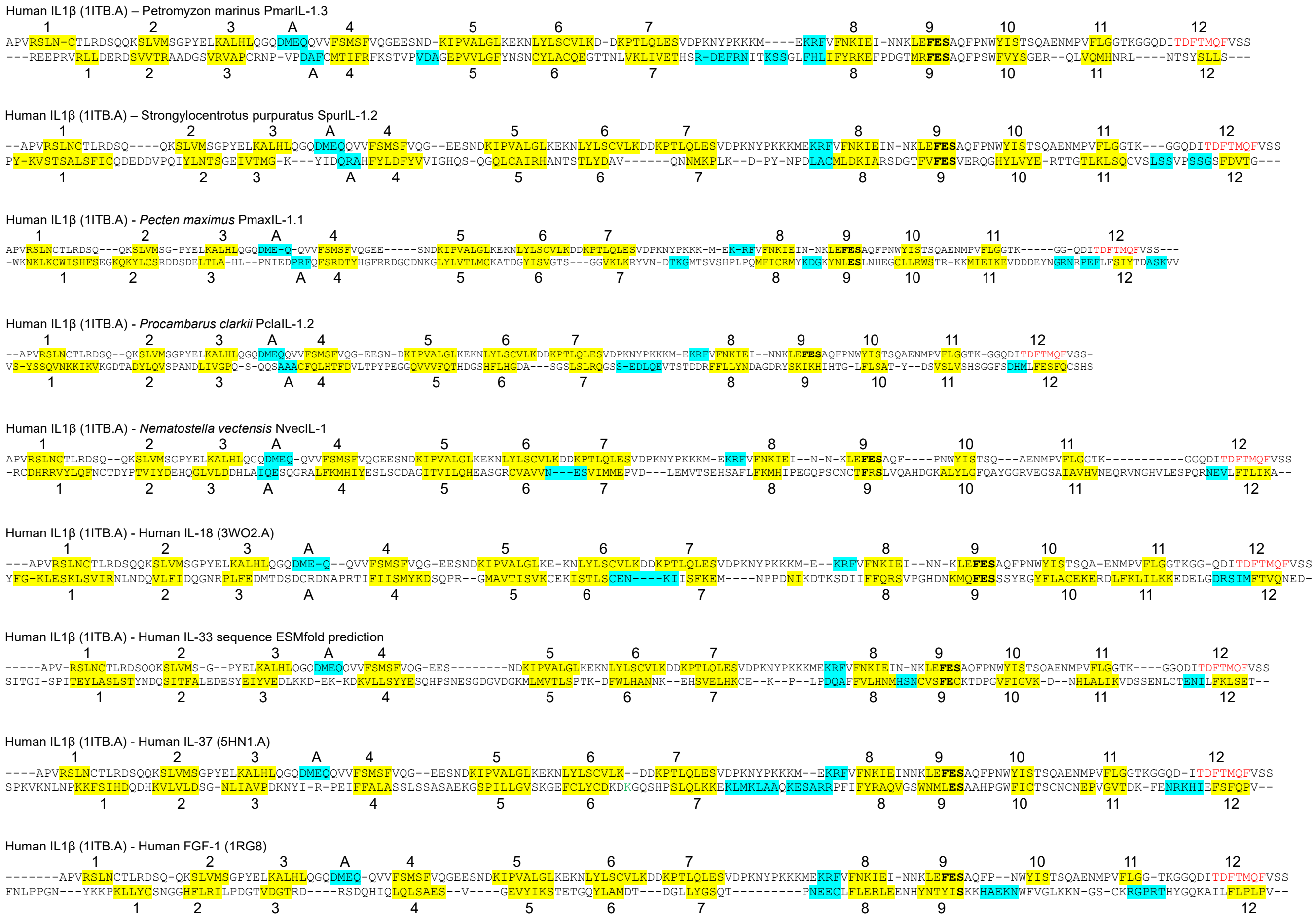


**Suppl. Figure S1. Evolution of IL-1**

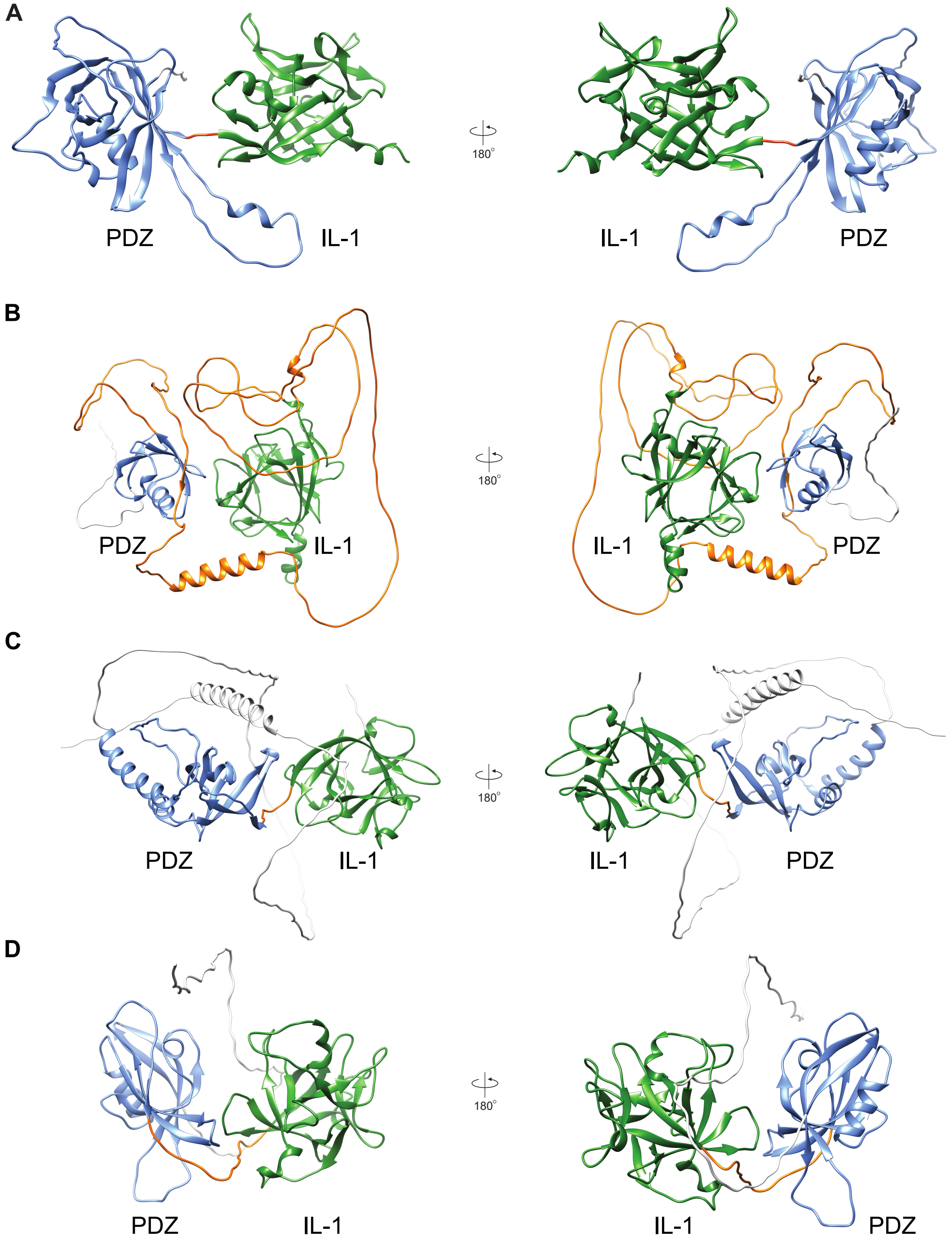


**Suppl. Figure S2. Evolution of IL-1**

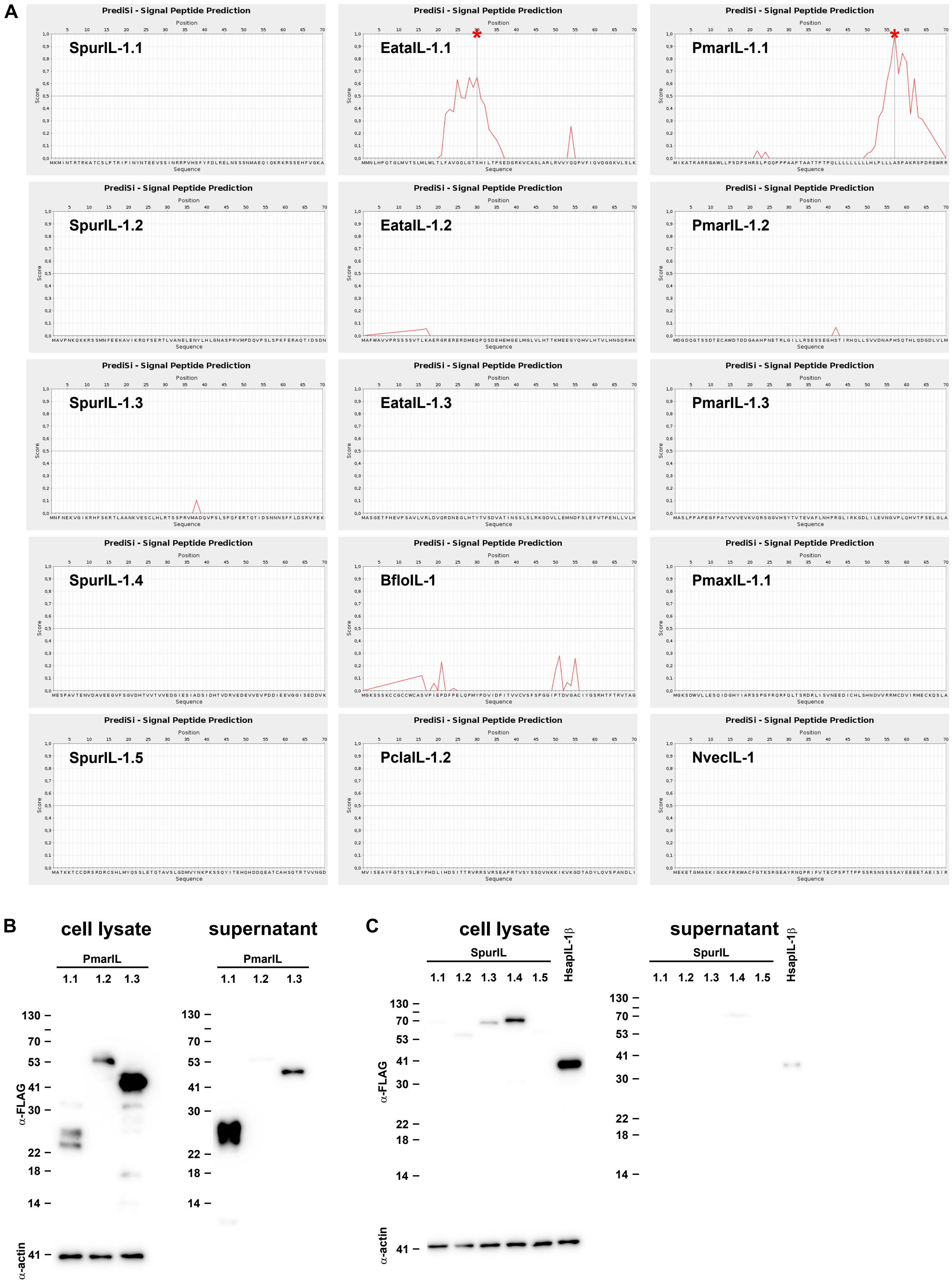


Suppl. Figure S3. Evolution of IL-1

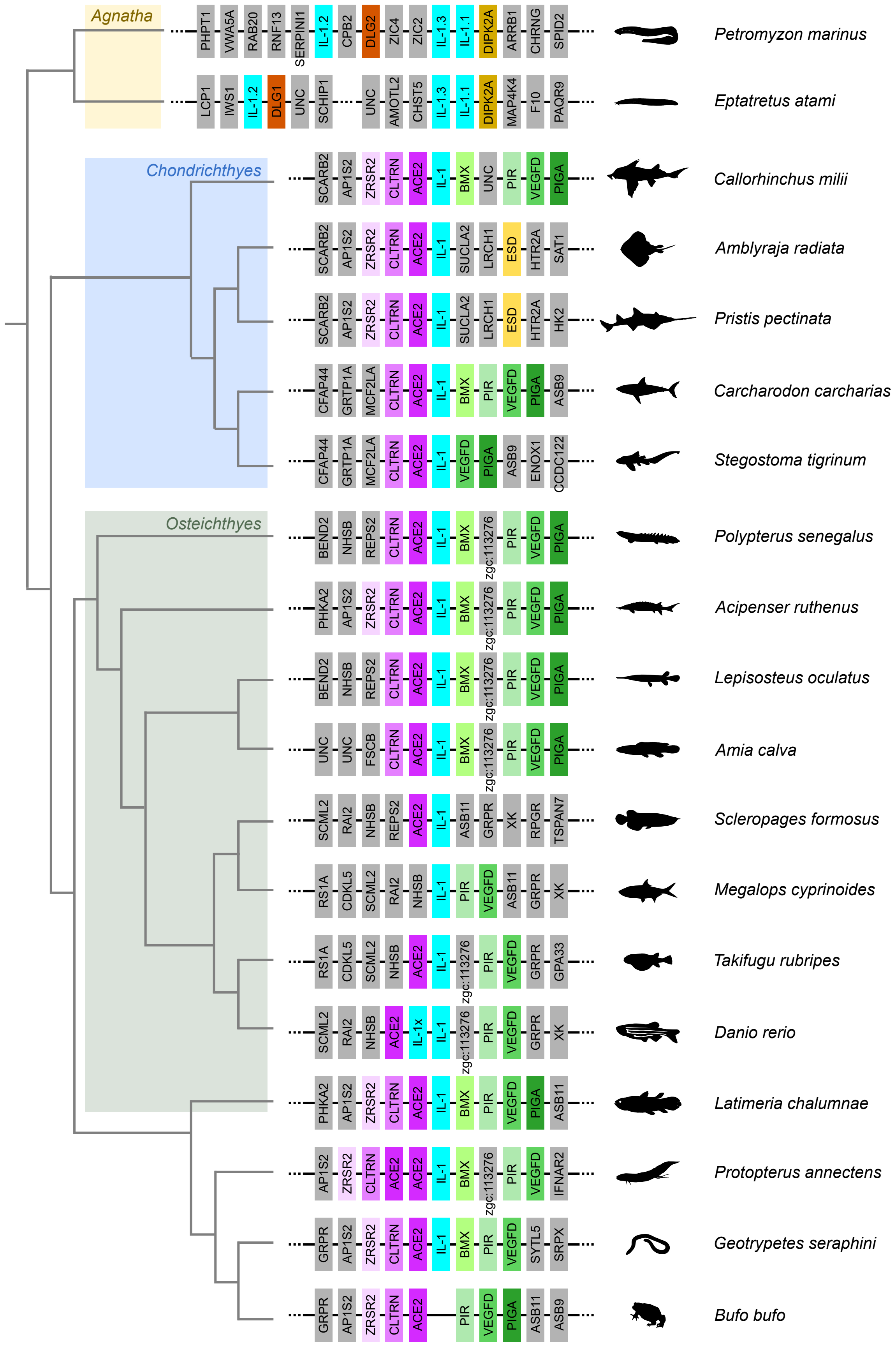


**Suppl. Figure S4. Evolution of IL-1**

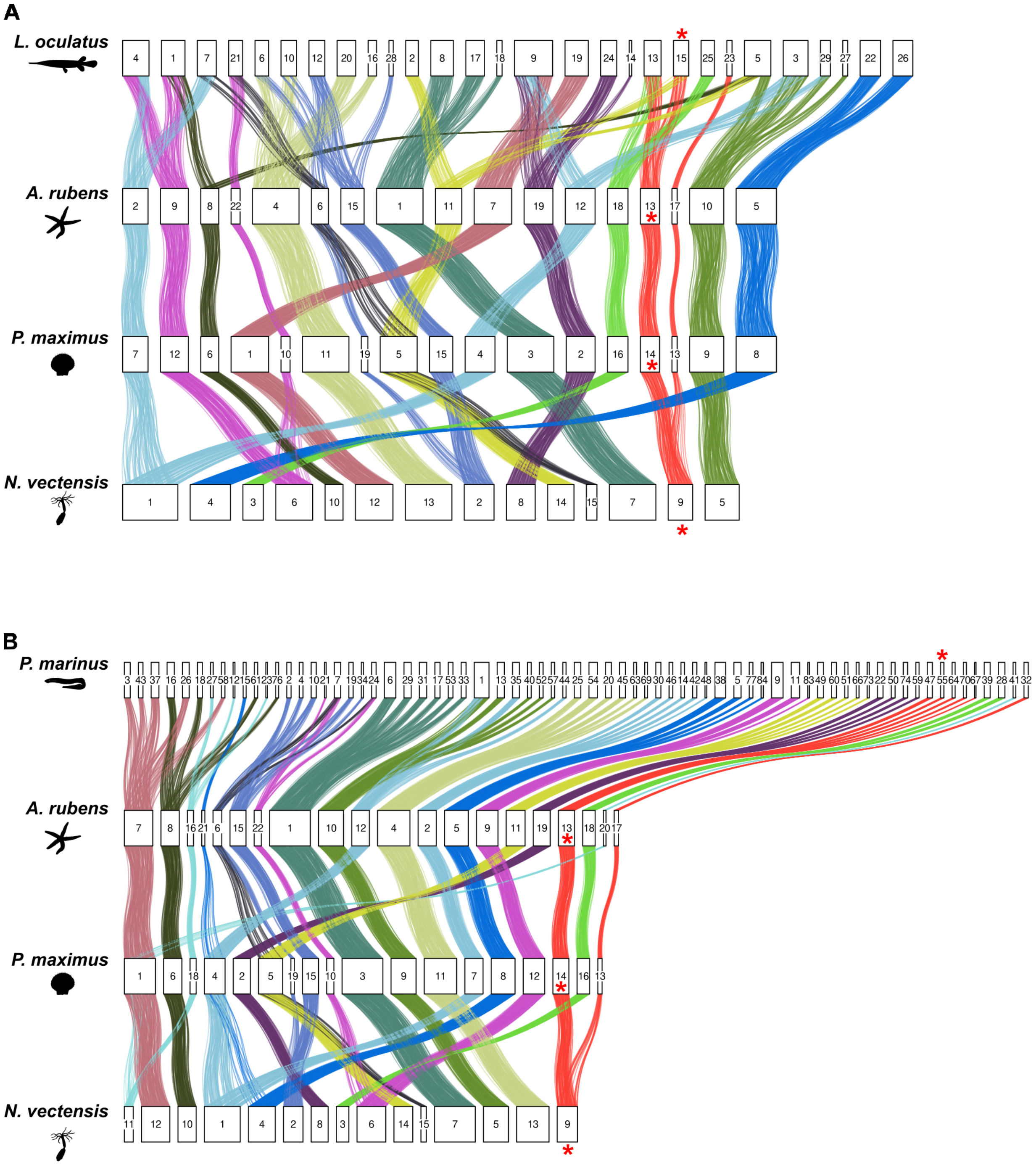


**Suppl. Figure S5. Evolution of IL-1**

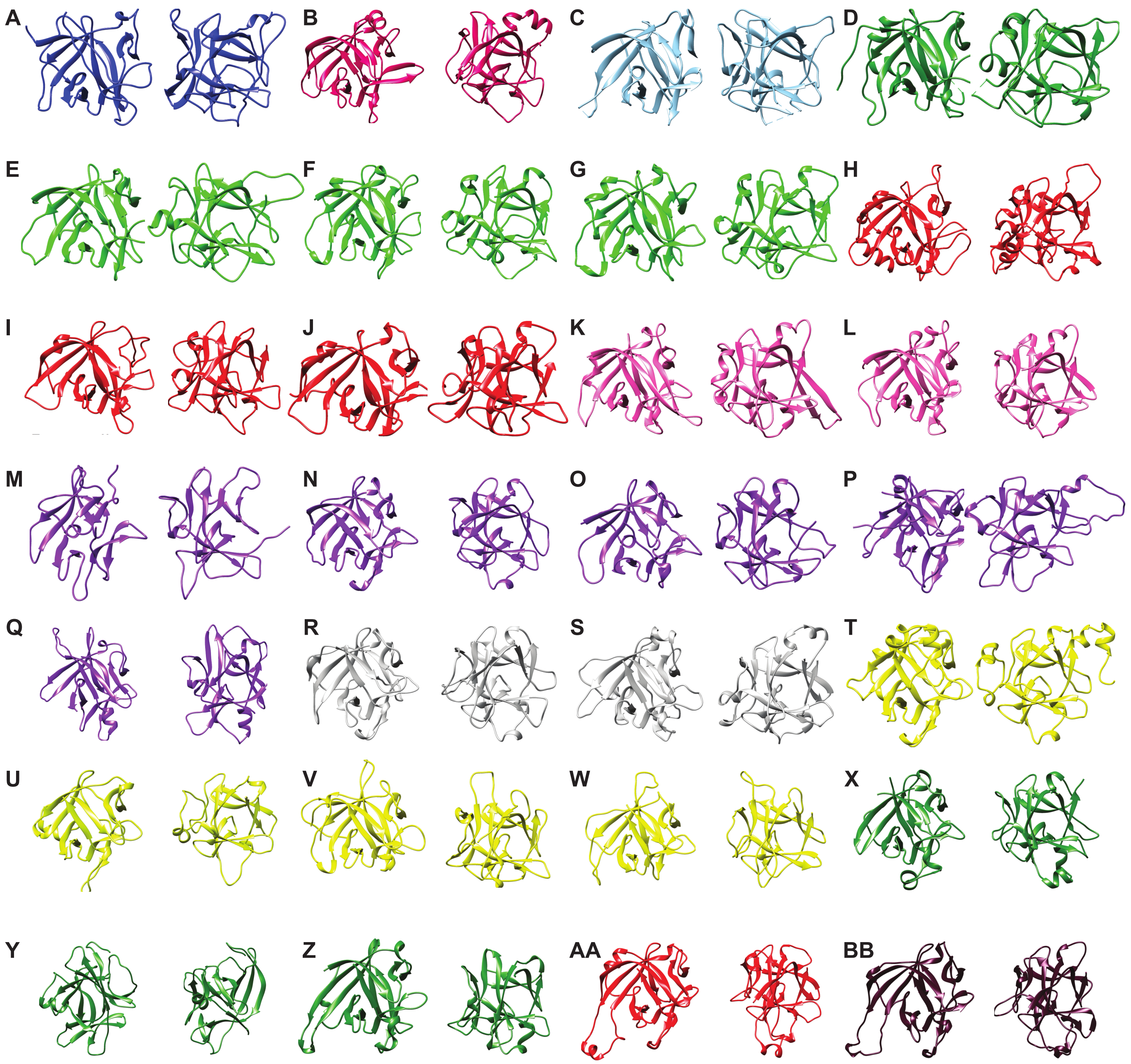


**Suppl. Figure S6. Evolution of IL-1**

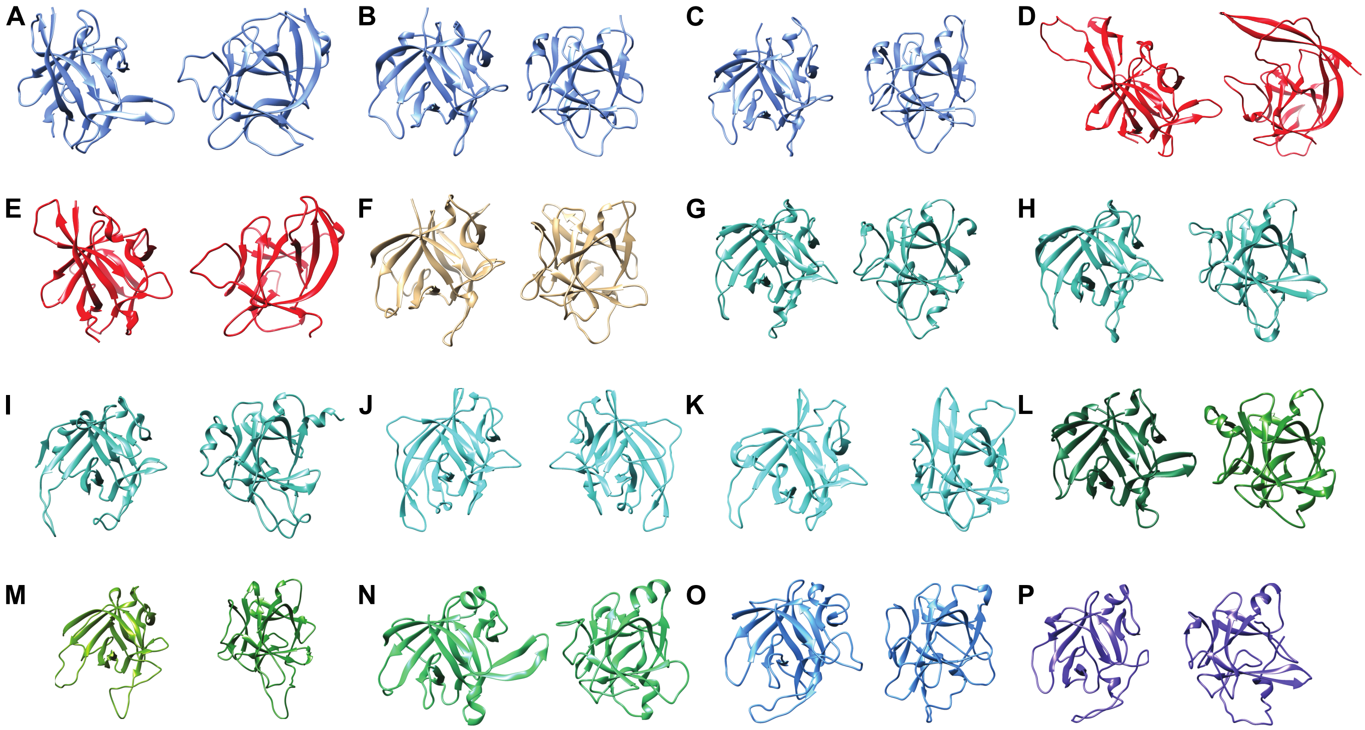


**Suppl. Figure S7. Evolution of IL-1**

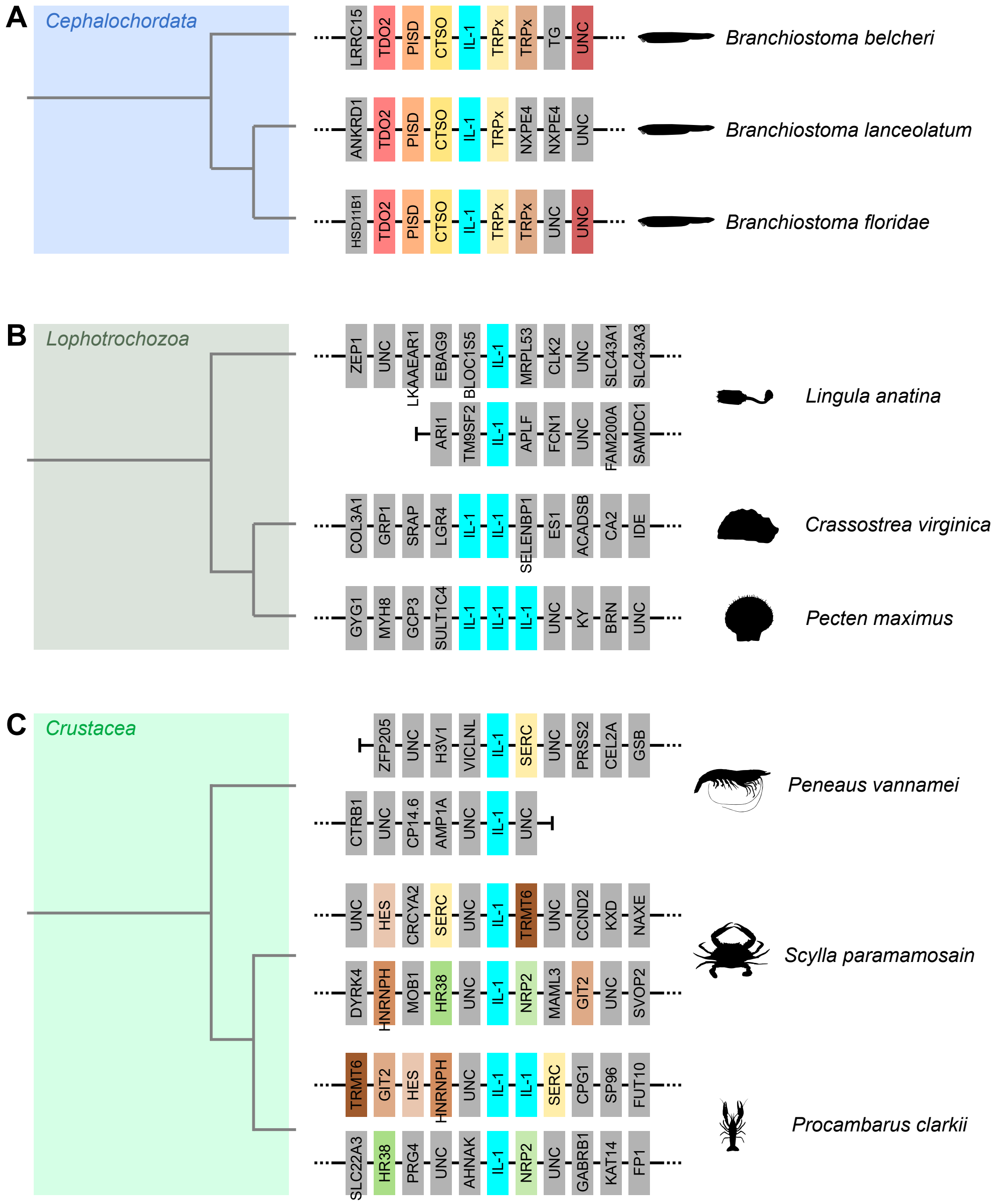


**Suppl. Figure S8. Evolution of IL-1**

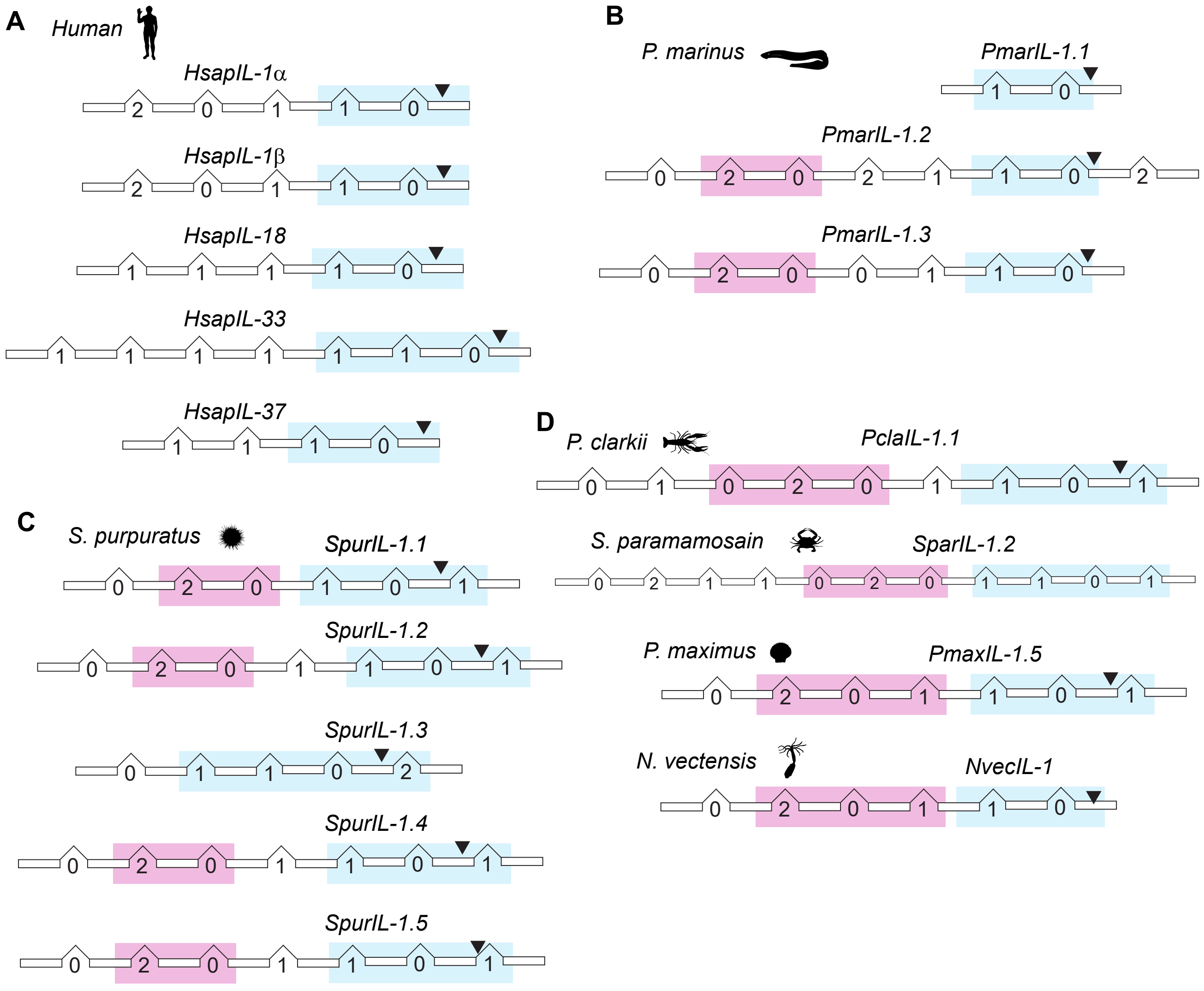


**Suppl. Figure S9. Evolution of IL-1**

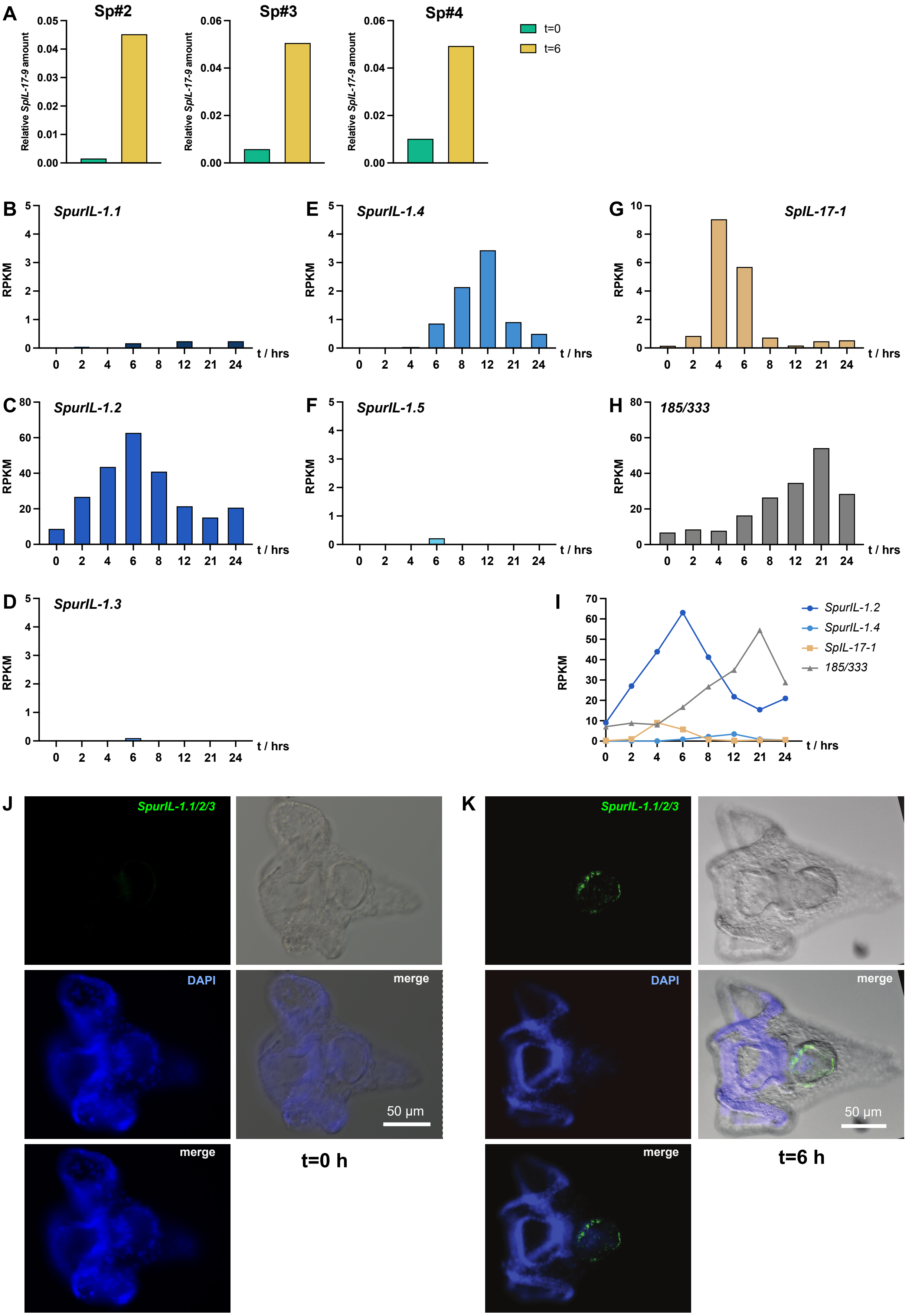


**Suppl. Figure S10. Evolution of IL-1**

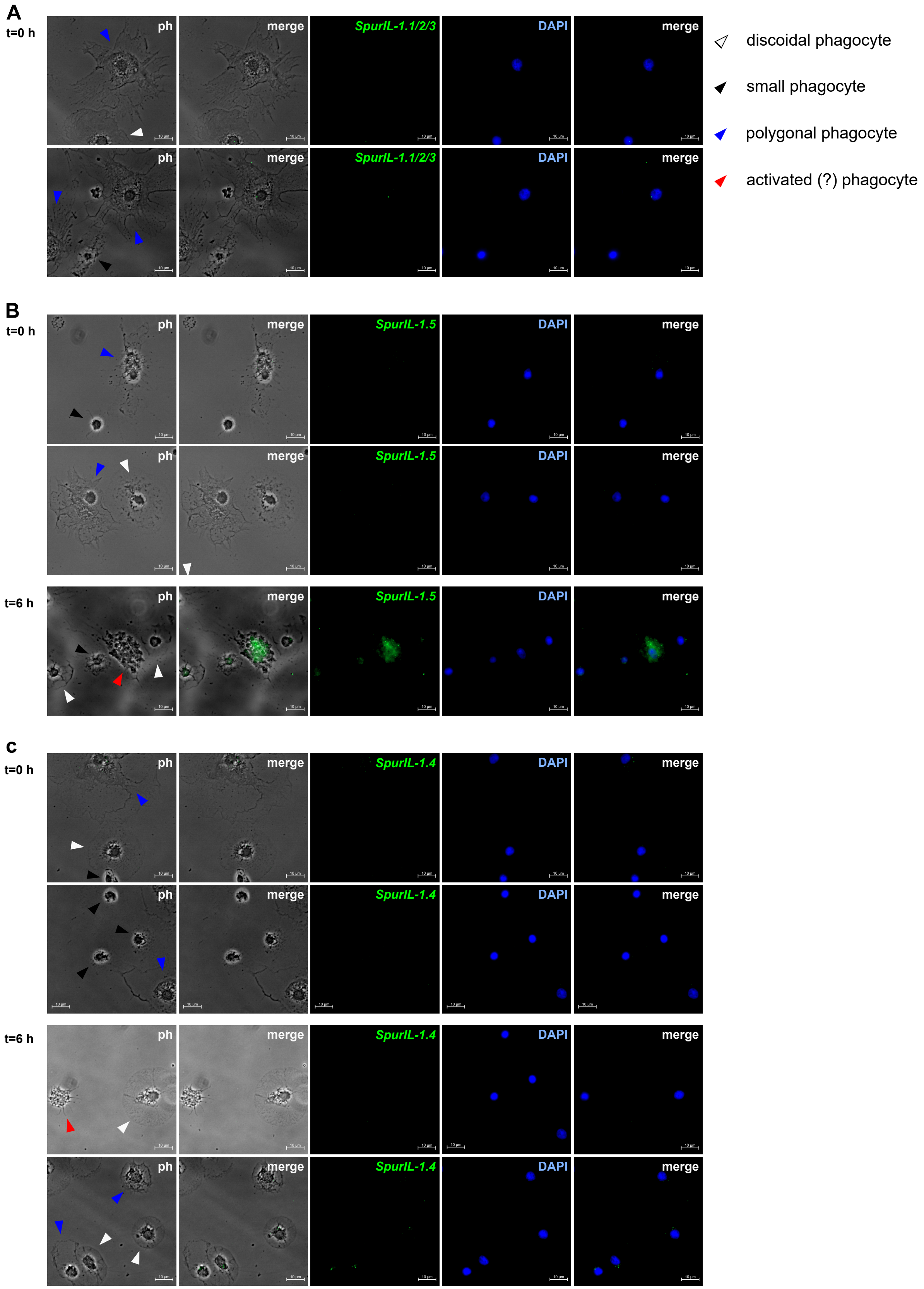


**Suppl. Figure S11. Evolution of IL-1**

### Supplemental Table 1 - List of genome assemblies

| | | | Common Name | Latin Name | Genome assembly | Source |
|---|---|---|---|---|---|---|
| Non-bilaterian r | Ctenophore | Ctenophore | Short-lobed comb jelly | Bolinopsis microptera | MBARI_Bmic_1.0 | GCF_026151205.1 |
| Non-bilaterian r | Porifera | demosponge | Great Barrier Reef sponge | Amphimedon queenslandica | v1.1 | GCF_000090795.2 |
| Non-bilaterian r | Cnidaria | Anthozoa | Green and Black Carnation | Dendronephthya gigantea | DenGig_1.0 | GCF_004324835.1 |
| Non-bilaterian r | Cnidaria | Anthozoa | Pulse coral | Xenia sp. Carnegie-2017 | XeniaSp_v1 | GCF_021976095.1 |
| Non-bilaterian r | Cnidaria | Anthozoa | Staghorn coran | Acropora millepora | Amil_v2.1 | GCF_013753865.1 |
| Non-bilaterian r | Cnidaria | Anthozoa | Cauliflower coral | Pocillopora damicornis | ASM370409v1 | GCF_003704095.1 |
| Non-bilaterian r | Cnidaria | Anthozoa | Mountainous star coral | Orbicella faveolata | ofav_dov_v1 | GCF_002042975.1 |
| Non-bilaterian r | Cnidaria | Anthozoa | Starlet sea anemone | Nematostella vectensis | jaNemVect1.1 | GCF_932526225.1 |
| Non-bilaterian r | Cnidaria | Anthozoa | Pale anemone | Exaiptasia diaphana | Aiptasia genome 1.1 | GCF_001417965.1 |
| Non-bilaterian r | Cnidaria | Anthozoa | Waratah anemone | Actinia tenebrosa | ASM960242v1 | GCF_009602425.1 |
| Protostome | Nematode | Chromadorea | | Caenorhabditis elegans | WBcel235 | GCF_000002985.6 |
| Protostome | Arthropod | Barnacle | Striped barnacle | Amphibalanus amphitrite | NRLGWU_Aamphi_draft | GCF_019059575.1 |
| Protostome | Arthropod | Isopod | Nosy pillbug | Armadillidium nasatum | CNRS_Arma_nasa_1.0 | GCA_009176605.1 |
| Protostome | Arthropod | Isopod | Common pillbug | Armadillidium vulgare | Arma_vul_BF2787 | GCA_004104545.1 |
| Protostome | Arthropod | Decapod | Green mud crab | Scylla paramamosain | ASM3559412v1 | GCF_035594125.1 |
| Protostome | Arthropod | Decapod | Red swamp crayfish | Procambarus clarkii | FALCON_Pclarkii_2.0 | GCF_040958095.1 |
| Protostome | Arthropod | Decapod | Whiteleg shrimp | Penaeus vannamei | ASM4276789v1 | GCF_042767895.1 |
| Protostome | Arthropod | Hexapod | Fruit fly | Drosophila melanogaster | Release 6 plus ISO1 MT | GCF_000001215.4 |
| Protostome | Annelid | Sedentaria | Bristle worm | Capitella teleta | Capca1 | GCA_000328365.1 |
| Protostome | Brachiopod | Lingulata | Duck lingula | Lingula anatina | LinAna2.0 | GCF_001039355.2 |
| Protostome | Mollusc | Bivalve | Great scallop | Pecten maximus | xPecMax1.1 | GCF_902652985.1 |
| Protostome | Mollusc | Bivalve | Eastern Oyster | Crassostrea virginica | C_virginica-3.0 | GCF_002022765.2 |
| Protostome | Mollusc | Bivalve | Pacific blue mussel | Mytilus trossulus | PNRI_Mtr1.1.1.hap1 | GCF_036588685.1 |
| Deuterostome | Invertebrate deuteros | Hemichordate | Hawaiian acorn worm | Ptychodera flava | AS_Pfla_20210202 | GCF_041260155.1 |
| Deuterostome | Invertebrate deuteros | Hemichordate | Virginia acorn worm | Saccoglossus kowalevskii | Skow_1.1 | GCF_000003605.2 |
| Deuterostome | Invertebrate deuteros | Echinoderm | Feather star | Anneissia japonica | ASM1163010v1 | GCF_011630105.1 |
| Deuterostome | Invertebrate deuteros | Echinoderm | Common seastar | Asterias rubens | eAstRub1.3 | GCF_902459465.1 |
| Deuterostome | Invertebrate deuteros | Echinoderm | Crown-of-thorns starfish | Acanthaster planci | OKI-Apl_1.0 | GCF_001949145.1 |
| Deuterostome | Invertebrate deuteros | Echinoderm | Bat star | Patiria miniata | Pmin_3.0 | GCF_015706575.1 |
| Deuterostome | Invertebrate deuteros | Echinoderm | Black sea cucumber | Holothuria leucospilota | CAS_HOLleu_V1 | GCA_029531755.1 |
| Deuterostome | Invertebrate deuteros | Echinoderm | Purple sea urchin | Strongylocentrotus purpuratu | Spur_5.0 | GCF_000002235.5 |
| Deuterostome | Invertebrate deuteros | Echinoderm | Painted sea urchin | Lytechinus pictus | Lp3.0 | GCF_037042905.1 |
| Deuterostome | Invertebrate deuteros | Echinoderm | Variegated sea urchin | Lytechinus variegatus | Lvar_3.0 | GCF_018143015.1 |
| Deuterostome | Invertebrate deuteros | Tunicate | Vase tunicate | Ciona intestinalis | KH | GCF_000224145.3 |
| Deuterostome | Invertebrate deuteros | Tunicate | Leathery sea squirt | Styela clava | ASM1312258v2 | GCF_013122585.1 |
| Deuterostome | Invertebrate deuteros | Tunicate | Star tunicate | Botryllus schlosseri | 356a-chromosome-assembly | GCA_000444245.1 |
| Deuterostome | Invertebrate deuteros | Cephalochordate | Florida amphioxus | Branchiostoma floridae | Bfl_VNyyK | GCF_000003815.2 |
| Deuterostome | Invertebrate deuteros | Cephalochordate | Common lancelet | Branchiostoma lanceolatum | klBraLanc5.hap2 | GCF_035083965.1 |
| Deuterostome | Invertebrate deuteros | Cephalochordate | Belcher's lancelet | Branchiostoma belcheri | Haploidv18h27 | GCF_001625305.1 |
| Deuterostome | Jawless vertebrate | Hagfish | Eptatretus atami | Eptatretus atami | Eptata_v1 | GCA_035128595.1 |
| Deuterostome | Jawless vertebrate | Hagfish | Eptatretus burgeri | Eptatretus burgeri | Eburgeri_v1 | GCA_024346535.1 |
| Deuterostome | Jawless vertebrate | Lamprey | Sea lamprey | Petromyzon marinus | kPetMar1 | GCF_010993605.1 |
| Deuterostome | Jawless vertebrate | Lamprey | Far Eastern brook lamprey | Lethenteron reissneri | ASM1570882v1 | GCF_015708825.1 |
| Deuterostome | Jawless vertebrate | Cartilaginous fishes | Elephant shark | Callorhinchus millii | IMCB_Cmil_1.0 | GCF_018977255.1 |
| Deuterostome | Jawed vertebrate | Cartilaginous fishes | Thorny skate | Amblyraja radiata | sAmbRad1.1.pri | GCF_010909765.2 |
| Deuterostome | Jawed vertebrate | Cartilaginous fishes | Smalltooth sawfish | Pristis pectinata | sPriPec2.1.pri | GCF_009764475.1 |
| Deuterostome | Jawed vertebrate | Cartilaginous fishes | Great white shark | Carcharodon carcharias | sCarCar2.pri | GCF_017639515.1 |
| Deuterostome | Jawed vertebrate | Cartilaginous fishes | Zebra shark | Stegostoma tigrinum | sSteTig4.hap1 | GCF_030684315.1 |
| Deuterostome | Jawed vertebrate | Cartilaginous fishes | Atlantic stingray | Hypanus sabinus | sHypSab1.hap1 | GCF_030144855.1 |
| Deuterostome | Jawed vertebrate | Cartilaginous fishes | Lesser devil ray | Mobula hypostoma | sMobHyp1.1 | GCF_963921235.1 |
| Deuterostome | Jawed vertebrate | Cartilaginous fishes | Epaulette shark | Hemiscyllium ocellatum | sHemOce1.pat.X.cur | GCF_020745735.1 |
| Deuterostome | Jawed vertebrate | Cartilaginous fishes | Whale shark | Rhincodon typus | sRhiTyp1.1 | GCF_021869965.1 |
| Deuterostome | Jawed vertebrate | Cartilaginous fishes | Smaller spotted catshark | Scyliorhinus canicula | sScyCan1.1 | GCF_902713615.1 |
| Deuterostome | Jawed vertebrate | Cartilaginous fishes | Little skate | Leucoraja erinacea | Leri_hhj_1 | GCF_028641065.1 |
| Deuterostome | Jawed vertebrate | Cartilaginous fishes | Smalled spotted catshark | Scyliorhinus torazame | Storazame_v1.0 | GCA_003427355.1 |
| Deuterostome | Jawed vertebrate | Osteichthyes | Gray bichir | Polypterus senegalus | ASM1683550v1 | GCF_016835505.1 |
| Deuterostome | Jawed vertebrate | Osteichthyes | Sterlet | Acipenser ruthenus | fAciRut3.2 maternal haplotype | GCF_902713425.1 |
| Deuterostome | Jawed vertebrate | Osteichthyes | Spotted gar | Lepisosteus oculatus | LepOcu1 | GCA_000242695.1 |
| Deuterostome | Jawed vertebrate | Osteichthyes | Bowfin | Amia calva | AmiCal1 | GCA_017591415.1 |
| Deuterostome | Jawed vertebrate | Osteichthyes | Asian arowana | Scleropages formosus | ASM162426v1 | GCA_001624265.1 |
| Deuterostome | Jawed vertebrate | Osteichthyes | Indo-Pacific tarpon | Megalops cyprinoides | fMegCyp1.pri | GCF_013368585.1 |
| Deuterostome | Jawed vertebrate | Osteichthyes | European eel | Anguilla anguilla | fAngAng1.pri | GCA_018320845.1 |

| | | | | | | |
|---|---|---|---|---|---|---|
| Deuterostome | Jawed vertebrate | Osteichthyes | Fugu | Takifugu rubripes | FUGU5 | GCA_000180615.2 |
| Deuterostome | Jawed vertebrate | Osteichthyes | Zebrafish | Danio rerio | GRCz11 | GCA_000002035.4 |
| Deuterostome | Jawed vertebrate | Osteichthyes | Southern platyfish | Xiphophorus maculatus | X_maculatus-5.0-male | GCF_002775205.1 |
| Deuterostome | Jawed vertebrate | Osteichthyes | Rainbow Trout | Oncorhynchus mykiss | USDA_OmykA_1.1 | GCF_013265735.2 |
| Deuterostome | Jawed vertebrate | Osteichthyes | Mexican tetra | Astyanax mexicanus | Astyanax_mexicanus-2.0 | GCA_000372685.2 |
| Deuterostome | Jawed vertebrate | Osteichthyes | Channel bull blenny | Cottoperca gobio | fCotGob3.1 | GCF_900634415.1 |
| Deuterostome | Jawed vertebrate | Osteichthyes | Old Calabar mormyrid | Paramormyrops kingsleyae | PKINGS_0.1 | GCA_002872115.1 |
| Deuterostome | Jawed vertebrate | Osteichthyes | Yellow perch | Perca flavescens | PFLA_1.0 | GCF_004354835.1 |
| Deuterostome | Jawed vertebrate | Osteichthyes | Amazon molly | Poecilia formosa | Poecilia_formosa-5.1.2 | GCA_000485575.1 |
| Deuterostome | Jawed vertebrate | Osteichthyes | Platy | Xiphophorus maculatus | X_maculatus-5.0-male | GCA_002775205.2 |
| Deuterostome | Jawed Vertebrate | Coelacanthiformes | Coelacanth | Latimeria chalumnae | fLatCha1.pri | GCF_037176945.1 |
| Deuterostome | Jawed Vertebrate | Ceratodontiformes | West African lungfish | Protopterus annectens | PAN1.0 | GCF_019279795.1 |
| Deuterostome | Jawed Vertebrate | Amphibian | Gaboon caecilian | Geotrypetes seraphini | aGeoSer1.1 | GCF_902459505.1 |
| Deuterostome | Jawed Vertebrate | Amphibian | | Microcaecilia unicolor | aMicUni1.1 | GCF_901765095.1 |
| Deuterostome | Jawed Vertebrate | Amphibian | Two-lined caecilian | Rhinatrema bivittatum | aRhiBiv1.1 | GCF_901001135.1 |
| Deuterostome | Jawed Vertebrate | Amphibian | Axolotl | Ambystoma mexicanum | UKY_AmexF1_1 | GCF_040938575.1 |
| Deuterostome | Jawed Vertebrate | Amphibian | Common toad | bufo bufo | aBufBuf1.1 | GCF_905171765.1 |
| Deuterostome | Jawed Vertebrate | Amphibian | Asiatic tod | Bufo gargarizans | ASM1485885v1 | GCF_014858855.1 |
| Deuterostome | Jawed Vertebrate | Amphibian | Hourglass treefrog | Dendropsophus ebraccatus | aDenEbr1.pat | GCF_027789765.1 |
| Deuterostome | Jawed Vertebrate | Reptilian | Bynoe's gecko | Heteronotia binoei | APGP_CSIRO_Hbin_v1 | GCF_032191835.1 |
| Deuterostome | Jawed Vertebrate | Reptilian | Green anole | Anolis carolinensis | rAnoCar3.1.pri | GCF_035594765.1 |
| Deuterostome | Jawed Vertebrate | Aves | Red junglefowl (chicken) | Gallus gallus | bGalGal1.mat.broiler.GRCg7b | GCF_016699485.2 |
| Deuterostome | Jawed Vertebrate | Mammal | House mouse | Mus musculus | GRCm39 | GCF_000001635.27 |
| Deuterostome | Jawed vertebrate | Mammal | Human | Homo sapiens | GRCh38.p14 | GCF_000001405.40 |

### Supplemental Table 2 - IL-1anc protein names and seqeunces

| Species | LOC | GenBank accession number | other accession number | original accession number | comments | common name | PDZ domain | amino acid sequence |
|---|---|---|---|---|---|---|---|---|
| Nematostella vectensis | LOC5506646 | XP_001627354.1 | | | | NvecIL-1 | y | |
| Exaiptasia diaphana (old name: Exaip | LOC110242051 | XXX_Acc#_TBD_XXX | | XP_020903649.2 | original XP incomplete, cloned and sequenced using RT-PCR | EpalIL-1 | y | MSSKFSKLFRKWSCLGGKTPSLEDEAQNNLRRN |
| Actinia tenebrosa | LOC116303115 | XP_031568442.1 | | | | AtenIL-1 | y | |
| Amphibalanus amphitrite | LOC122367971 | XP_043197481.1 | | | | AampIL-1 | y | |
| Armadillidium nasatum | Anas_05996 | KAB7502784.1 | | | | AnasIL-1 | y | |
| Armadillidium vulgare | Avbf_04417 | RXG58008.1 | | | | AvulIL-1 | y | |
| Scylla paramamosain | LOC135103869 | XP_063866844.1 | | | | SparIL-1.1 | y | |
| | LOC135103538 | XP_063866017.1 | | | | SparIL-1.2 | n | |
| Procambarus clarkii | LOC123746562 | XP_045584109.1 | | | | PclaIL-1.1 | y | |
| | LOC123746547 | XP_069174462.1 | | | | PclaIL-1.2 | n | |
| | LOC123746438 | XP_069174463.1 | | | | PclaIL-1.3 | n | |
| Peneaus vannamei | LOC113809227 | XP_027216559.1 | | | | PvanIL-1.1 | y | |
| | LOC113801232 | XP_027207847.1 | | | | PvanIL-1.2 | n | |
| Lingula anatina | LOC106166605 | XP_013400692.1 | | | | LanaIL-1.1 | y | |
| | LOC106154450 | XP_013384251.1 | | | | LanaIL-1.2 | y | |
| Pecten maximus | LOC117342263 | XP_033760242.1 | | | | PmaxIL-1.1 | y | |
| | LOC117342765 | XP_033760898.1 | | | | PmaxIL-1.2 | y | |
| | LOC117342269 | XP_033760250.1 | | | | PmaxIL-1.3 | y | |
| Crassostrea virginica | LOC111101316 | XP_022289471.1 | | | | CvirIL-1.1 | y | |
| | LOC111100639 | XP_022288429.1 | | | | CvirIL-1.2 | y | |
| Mytilus trossulus | LOC134709531 | XP_063425759.1 | | | | MtroIL-1.1 | y | |
| | LOC134709801 | XP_063426011.1 | | | | MtroIL-1.2 | y | |
| Ptychodera flava | LOC139118898 | XP_070538562.1 | | | | PflaIL-1 | y | |
| Saccoglossus kowalevskii | LOC100378855 | XP_006813206.1 | | | | SkowIL-1.1 | y | |
| | LOC100379003 | XP_006813207.1 | | | | SkowIL-1.2 | y | |
| Anneissia japanoica | LOC117117896 | XP_033118264.1 | | | | AjapIL-1.1 | n | |
| | LOC117117911 | XP_033118281.1 | | | | AjapIL-1.2 | y | |
| | LOC117106586 | XP_033103862.1 | | | | AjapIL-1.3 | y | |
| Asterias rubens | LOC117298237 | XP_033637260.1 | | | | ArubIL-1.1 | y | |
| | LOC117295996 | XP_033634717.1 | | | | ArubIL-1.2 | y | |
| | LOC117293103 | XP_033631221.1 | | | | ArubIL-1.3 | y | |
| | LOC117295764 | XP_033634398.1 | | | | ArubIL-1.4 | y | |
| Acanthaster planci | LOC110981036 | XP_022093886.1 | | | | AplaIL-1.1 | y | |
| | LOC110985595 | XP_022102418.1 | | | | AplaIL-1.2 | y | |
| | LOC110985596 | XP_022102420.1 | | | | AplaIL-1.3 | y | |
| | LOC110983422 | XP_022098364.1 | | | | AplaIL-1.4 | y | |
| | LOC110982626 | XP_022096875.1 | | | | AplaIL-1.5 | y | |
| | LOC110982596 | XP_022096825.1 | | | | AplaIL-1.6 | y | |
| | LOC110982587 | XP_022096795.1 | | | | AplaIL-1.7 | y | |
| Patiria miniata | LOC119723747 | XP_038050513.1 | | | | PminIL-1.1 | y | |
| | LOC119723861 | XP_038050670.1 | | | | PminIL-1.2 | y | |
| | LOC119723835 | XP_038050642.1 | | | | PminIL-1.3 | y | |
| | LOC119737400 | XP_038067657.1 | | | two separate IL-1 genes in one LOC | PminIL-1.4 | y | |
| | LOC119737400 | XP_038067653.1 | | | two separate IL-1 genes in one LOC | PminIL-1.5 | y | |
| | LOC119720123 | XP_038045603.1 | | | | PminIL-1.6 | y | |
| | LOC119719369 | XP_038044726.1 | | | | PminIL-1.7 | y | |
| | LOC119720121 | XP_038045599.1 | | | | PminIL-1.8 | y | |
| | LOC119737842 | XP_038068394.1 | | | | PminIL-1.9 | y | |
| | LOC119720977 | XP_038046770.1 | | | | PminIL-1.10 | y | |
| Holothuria leucospilota | HOLleu_27639 | KAJ8031043.1 | | | | HLeuIL-1.1 | y | |

| | | | | | | | | |
|---|---|---|---|---|---|---|---|---|
| | HOLleu_27637 | KAJ8031042.1 | | | | HLeuIL-1.2 | y | |
| Strongylocentrotus purpuratus | LOC105446168 | XXX_Acc#_TBD_XXX | | XP_030837460.1 | cloned | SpurIL-1.1 | y | MKMINTRTRKATCSLPTRIPINYNTEEVSSINRRPV |
| | LOC583539 | XXX_Acc#_TBD_XXX | | XP_800233.2 | cloned | SpurIL-1.2 | y | MAVPNKQKKRSSMNFEEKAVIKRQFSERTLVANI |
| | LOC105446162 | XXX_Acc#_TBD_XXX | | XP_030838393.1 | cloned, original annotation<br>lacked 5'end of gene | SpurIL-1.3 | n | MNFNEKVGIKRHFSKRTLAANKVESCLHLRTSSF |
| | LOC105446163 | XXX_Acc#_TBD_XXX | | XP_011680905.2 | cloned | SpurIL-1.4 | y | MESPAVTENVDAVEEGVFSGVDHTVVTVVEDGIE |
| | LOC115918145+<br>LOC105441282+<br>LOC115918179 | XXX_Acc#_TBD_XXX | | XP_030838363.1+<br>XP_030837393.1 | cloned, original annotation was<br>split into three separate genes | SpurIL-1.5 | y | MATKKTCCDRSRDRCSHLMYQSSLETQTAVSLG |
| Lytechinus pictus | LOC129278821 | XP_054770931.2 | | | | LpicIL-1.1 | y | |
| | LOC129278820 | XP_054770929.2 | | | | LpicIL-1.2 | y | |
| | LOC129260727 | XP_063970477.1 | | | | LpicIL-1.3 | y | |
| | LOC129260728 | XP_054754655.2 | | | | LpicIL-1.4 | y | |
| | LOC129279149 | XP_054771227.2 | | | | LpicIL-1.5 | y | |
| Lytechinus variegatus | LOC121429905 | XP_041483100.1 | | | | LvarIL-1.1 | y | |
| | LOC121429914 | XP_041483110.1 | | | | LvarIL-1.2 | y | |
| | LOC121416031 | XP_041465404.1 | | | | LvarIL-1.3 | y | |
| | LOC121415439 | XP_041464575.1 | | | | LvarIL-1.4 | y | |
| | LOC121430357 | XP_041483571.1 | | | | LvarIL-1.5 | y | |
| Branchiostoma floridae | LOC118419816 | XP_035682314.1 | | | | BfloIL-1 | y | |
| Branchiostoma lanceolatum | LOC136437018 | XP_066287540.1 | | | | BlanIL-1 | y | |
| Branchiostoma belcheri | LOC109487726 | XP_019647342.1 | | | | BbelIL-1 | y | |
| Eptatretus atami | | | | | manual annotation, signal peptide | EataIL-1.1 | n | MMNLHPQTGLMVTSLMLWLTLFAVGQLGTSHIL |
| | | | EA83902 in Pata_ah2p.pep.fa at<br>https://zenodo.org/records/10227719 | | | EataIL-1.2 | y | |
| | | | EA83591 in Pata_ah2p.pep.fa at<br>https://zenodo.org/records/10227719 | | | EataIL-1.3 | y | |
| Eptatretus burgeri | ENSEBUG00000011846.1 | | ENSEBUP00000019000.1<br>(Eburgeri_3.2 at Ensembl) | | signal peptide | EburIL-1.1 | n | |
| | ENSEBUG00000012019.1 | | | ENSEBUP00000019307.1<br>(Eburgeri_3.2 at Ensembl) | partial, manual annotation,<br>5'end not annotated | EburIL-1.2 | y | EENEMGELMGLVLHTTKMEEGYQHVLHTVLHN |
| | ENSEBUG00000001301.1 | | ENSEBUP00000001554<br>(Eburgeri_3.2 at Ensembl) | | | EburIL-1.3 | y | |
| Petromyzon marinus | LOC116954500 | XP_032830942.1 | | | signal peptide | PmarIL-1.1 | n | |
| | LOC116954507 | XP_032830949.1 | | | | PmarIL-1.2 | y | |
| | LOC116954430 | XP_032830829.1 | | | | PmarIL-1.3 | y | |
| Lethenteron reissneri | LOC133357515 | XP_061431465.1 | | | signal peptide | LreiIL-1.1 | n | |
| | IL36RN | XP_061431479.1 | | | partial, 5' end of gene not annotated | LreiIL-1.2 | y | |
| | LOC133357510 | XP_061431458.1 | | | | LreiIL-1.3 | y | |
| Callorhinchus milii | LOC103177481 | XP_007889844.1 | | | | CmilIL-1anc | y | |
| Amblyraja radiata | LOC116980568 | XP_032888811.1 | | | | AradIL-1anc | y | |
| Pristis pectinata | LOC127569811 | XP_051870694.1 | | | | PpecIL-1anc | y | |
| Carcharodon carcharias | LOC121290734 | XP_041067518.1 | | | | CcarIL-1anc | y | |
| Stegostoma tigrinum | LOC125457239 | XP_048397147.1 | | | | StigIL-1anc | y | |
| Polypterus senegalus | LOC120523605 | XP_039600982.1 | | | | PsenIL-1anc | y | |
| Acipenser ruthenus | LOC117406507 | XP_033866477.1 | | | | ArutIL-1anc | y | |
| Lepisosteus oculatus | LOC102690436 | XP_015219154.1 | | | | LoculIL-1anc | y | |
| Amia calva | LOC136750781 | XP_066561938.1 | | | | AcalIL-1anc | y | |
| Scleropages formosus | LOC108920476 | XP_018584753.1 | | | | SforIL-1anc | y | |
| Megalops cyprinoides | il1fma | XP_036401909.1 | | | | McypIL-1anc | y | |
| Anguilla anguilla | il1fma | XP_035263746.1 | | | | AangIL-1anc | y | |
| Takifugu rubripes | nil1fm | XP_011608335.1 | | | | TrubIL-1anc | y | |

| | | | | | | | |
|---|---|---|---|---|---|---|---|
| Danio rerio | il1fma | NP_001277347.1 | | | DrerIL-1anc.1 | y | |
| | LOC100150092 | NP_001269007.1 | | | DrerIL-1anc.2 | n | |
| Xiphophorus maculatus | LOC102218441 | XP_023188682.1 | | | XmacIL-1anc | y | |
| Latimeria chalumnae | | | XP_005997916.1 | not annotated (but present) in current genome version | LchaIL-1anc | y | |
| Protopterus annectens | LOC122803295 | XP_043928801.1 | | | PannIL-1anc | y | |
| Geotrypetes seraphini | LOC117362118 | XP_033803848.1 | | | GserIL-1anc | y | |
| Rhinatrema bivittatum | | | | partial, manual annotation, 5'end not annotated | RbivIL-1anc | y | MVHKEDASLPQEEAALSVHNGTCLECPGIEGED |

**Supplemental Table 3**

### Primers for RT-PCR expression analysis

| Target | Primer | Sequence | Purpose |
|---|---|---|---|
| *SpIL-1b1* | IL1B1-QF3 | CAGCATGAAGCTGATGGAGG | qPCR F |
| | IL1B1-QR3 | AGTGTTCCTGCAGCTCTGTC | qPCR R |
| *SpIL-1b2* | IL1B2-QF3 | TTCGTGTTCGAGTCAGTGGAG | qPCR F |
| | IL1B2-QR3 | ACCGAACTTAGACTCACGCA | qPCR R |
| *SpIL-1b3* | IL1B3-QF3 | GTCCTGATCAAGCATGTCTGC | qPCR F |
| | IL1B3-QR3 | TAAGTGCTCCTGCATCCCTG | qPCR R |
| *SpIL-1b4* | IL1B4-QF3 | TGACGGATCATCTGTGTTCCA | qPCR F |
| | IL1B4-QR3 | CGAATCGTCCTGACATCGGT | qPCR R |
| *SpIL-1b5* | IL1B5-QF3 | GGATCGTATCCACACTCCTGA | qPCR F |
| | IL1B5-QR3 | CGGGCCTGATCCATAGAACG | qPCR F |
| *SpIL-17 IX* | IL17-IX qPCRF | CAATCAGGAGCCTCTCRAGT | qPCR F |
| | IL17-IX qPCRR | AGGGTTAATACAATCACGGCAC | qPCR R |
| *18S (S.purp.)* | 18SF | CAGGGTTCGATTCCGTAGAG | qPCR F |
| | 18SF | CCTCCAGTGGATCCTCGTTA | qPCR R |
| *PmIL-1.1* | qIL1.1-F | AGTGCTATCCCGACGCTCCT | qPCR F |
| | qIL1.1-R | ACCGAAACCCAGCGACACGA | qPCR R |
| *PmIL-1.2* | qIL1.2-F | AGTCGCTACCTGTGCTGCTCA | qPCR F |
| | qIL1.2-R | TGGGTCAGCGGCTCTTCCTT | qPCR R |
| *PmIL-1.3* | qIL1.3-F | CGGTGGTGCTGGGCTTTTACA | qPCR F |
| | qIL1.3-R | TCGTCCCTGCTGTGAGTCTCCA | qPCR R |
| *PmIL-8* | qIL8-F | CCATCCCAAGCATTTCCAGACA | qPCR F |
| | qIL8-R | GGTGTCTGAGCCCCGTCCAAAA | qPCR R |
| *PmTNFα* | qTNF-F | CGAGAGGGGCTTCACGGTTGAT | qPCR F |
| | qTNF-R | TGAGATTGGTGAGGTCCTGGCG | qPCR R |
| *PmActin* | qbActin-F1 | GCCAACCGTGAAAAGATGACA | qPCR F |
| | qbActin-R1 | GGATGGCGACGTACATTGC | qPCR R |

**Supplemental Table 4**

### Primers for cloning SpIL-1s, PmIL-1s, and caspases

| Target | Primer | Sequence | Purpose |
|---|---|---|---|
| *SpIL1b1* | SpIL1b1F | ATGAAAATGATAAATACCAGGACTCGC | Clone into pJET1.2 |
| | SpIL1b1R | TACACGAGGGATACCGAAAGAG | |
| | SpIL1b1_seq_F2 | ATTCAGCAAGGTACAGGTGG | Sequencing (internal) |
| | SpIL1b1_SpeI_F | GGACTAGTACCATGAAAATGATAAATACCAG | Clone into pCMV-FLAG-IRES-GFP |
| | SpIL1b1_SalI_R | ACGCGTCGACTACACGAGGGATACCGAAAG | |
| *SpIL1b2* | SpIL1b2F | ATGGCTGTACCAAACAAGCAG | Clone into pJET1.2 |
| | SpIL1b2R | CACCCTTGAGGATTTGACACCAG | |
| | SpIL1b2_SpeI_F | GGACTAGTaccATGGCTGTACCAAACAAGCA | Clone into pCMV-FLAG-IRES-GFP |
| | SpIL1b2_SalI_R | ACGCGTCGACCACCCTTGAGGATTTGACAC | |
| *SpIL1b3* | SpIL1b3F | CAGCAGTTCCAATATTGCTATACC | Clone into pJET1.2 |
| | SpIL1b3R | AGTCCGGCTCGTACATACAC | |
| | SpIL1b3_BamHI_F | CGGGATCCaccATGAATTTTAATGAGAAGGT | Clone into pCMV-FLAG-IRES-GFP |
| | SpIL1b3_SalI_R | ACGCGTCGACCACATCGGACTTAACGGGTG | |
| *SpIL1b4* | SpIL1b4F | ATGGAGTCTCCAGCTGTAACAG | Clone into pJET1.2 |
| | SpIL1b4R | ATGTGCTCCATTCACATGTC | |
| | SpIL1b4_BamHI_F | CGGGATCCaccATGGAGTCTCCAGCTGTAAC | Clone into pCMV-FLAG-IRES-GFP |
| | SpIL1b4_XhoI_R | CCGCTCGAGATGTGCTCCATTCACATGTC | |
| SpIL1b5 | SpIL1b5F | CATTACTTGTTGTCATGGCGA | Clone into pJET1.2 |
| | SpIL1b5R | ATGGCTAAACATCGTAAGTTCTG | |
| | SpIL1b5_BamHI_F | CGGGATCCaccATGGCGACAAAGAAGACATG | Clone into pCMV-FLAG-IRES-GFP |
| | SpIL1b5_SalI_R | ACGCGTCGACAACATCGTAAGTTCTGAAGA | |
| *PmIL1.1* | PmarIL1.1_SpeI_F | GGACTAGTACCATGATCAAGGCCACCAGGGC | Clone into pCMV-FLAG-IRES-GFP |
| | PmarIL1.1_XhoI_R | CCGCTCGAGACCTCTCCTCACGGCTGGCACAA | |
| *PmIL1.2* | PmarIL1.2_SpeI_F | GGACTAGTACCATGGACGGCGATCAGGGCAC | Clone into pCMV-FLAG-IRES-GFP |
| | PmarIL1.2_XhoI_R | CCGCTCGAGACCAAACCTCTGTTTGGTGAAGC | |
| *HsCASP1* | HsCASP1_F1 | ATGGCCGACAAGGTCCTGAA | Clone into pCDNA3.1 V5/His |
| | HsCASP1_R1 | ATGTCCTGGGAAGAGGTAGA | |
| *PmCasp1* | PmarCASP1_F | accATGAGCGGCGACCCCAATCT | Clone into pCDNA3.1 V5/His |
| | PmarCASP1_R | GAAGCCAGGGAACAAGTAGA | |

**Supplemental Table 5**

### Primers for site-directed mutagenesis

| Target | Primer | Sequence | Purpose |
|---|---|---|---|
| *SpIL1b3* | SpIL1b3_D181A_F | CGAGAAGGAGGGGTGTgccGAACCCGATGGACTCT | D181A mutant |
| | SpIL1b3_D181A_R | AGAGTCCATCGGGTTCggcACACCCCTCCTTCTCG | |
| | SpIL1b3_D184A_F | GGGGTGTGACGAACCCgctGGACTCTTTGGTAGAA | D184A mutant |
| | SpIL1b3_D184A_R | TTCTACCAAAGAGTCCagcGGGTTCGTCACACCCC | |
| | SpIL1b3_D212A_F | CCTGAGAAAATATGGAgcaAAGGTTCATCTCGAGA | D212A mutant |
| | SpIL1b3_D212A_R | TCTCGAGATGAACCTTtgcTCCATATTTTCTCAGG | |
| | SpIL1b3_D226A_F | GATACGGGTCACGAAAgcaGATAAAGATGTGGTGT | D226A mutant |
| | SpIL1b3_D226A_R | ACACCACATCTTTATCtgcTTTCGTGACCCGTATC | |
| | SpIL1b3_D227A_F | ACGGGTCACGAAAGATgcaAAAGATGTGGTGTGCG | D227A mutant |
| | SpIL1b3_D227A_R | CGCACACCACATCTTTtgcATCTTTCGTGACCCGT | |
| | SpIL1b3_D229A_F | CACGAAAGATGATAAAgcaGTGGTGTGCGAAGGGC | D229A mutant |
| | SpIL1b3_D229A_R | GCCCTTCGCACACCACtgcTTTATCATCTTTCGTG | |
| *HsCasp1* | HsCASP1_C285A_F | GATCATCATCCAGGCCgcaCGTGGTGACAGCCCTG | C285A mutant |
| | HsCASP1_C285A_R | CAGGGCTGTCACCACGtgcGGCCTGGATGATGATC | |
